# The speed of GTP hydrolysis determines GTP cap size and controls microtubule stability

**DOI:** 10.1101/779108

**Authors:** Johanna Roostalu, Claire Thomas, Nicholas I. Cade, Simone Kunzelmann, Ian A. Taylor, Thomas Surrey

## Abstract

Microtubules are bistable cytoskeletal polymers whose function depends on their property to switch between states of growth and shrinkage ^1^. Growing microtubules are thought to be stabilized by a GTP cap at their ends ^2-5^. The nature of this cap, however, is still poorly understood. How GTP hydrolysis determines the properties of the GTP cap and hence microtubule stability is unclear. End Binding proteins (EBs) recruit a diverse range of regulators of microtubule function to growing microtubule ends ^6^. Whether these regulatory platforms at growing microtubule ends are identical to the GTP cap is not known. Using mutated human tubulin with blocked GTP hydrolysis, we demonstrate in microscopy-based *in vitro* reconstitutions that EB proteins bind with high affinity to the GTP conformation of microtubules. Slowing-down GTP hydrolysis leads to extended GTP caps and consequently hyper-stable microtubules. Single molecule experiments reveal that the microtubule conformation gradually changes in the cap as GTP is hydrolyzed. These results demonstrate the critical importance of the kinetics of GTP hydrolysis for microtubule stability; and establish that the GTP cap coincides with the EB-binding regulatory hub that modulates microtubule cytoskeleton function in cells.

## MAIN TEXT

The dynamic nature of microtubules is critical for their function in cells. Microtubules polymerize by the addition of GTP-bound α/β-tubulin heterodimers to their ends. Tubulin incorporation induces GTP hydrolysis which destabilizes the microtubule wall. A delay in hydrolysis of unknown duration is thought to produce a ‘GTP cap’, a protective end structure formed of GTP-tubulins that is critical for microtubule stability ^1-5^. The loss of this cap is thought to expose an unstable GDP lattice and trigger depolymerization, resulting in dynamic instability of microtubule growth ^1^. How the biochemistry of GTP hydrolysis translates into conformational changes in the growing microtubule end, and thereby determines microtubule stability, is not understood ^2, 7^.

Non-hydrolysable GTP analogues have been used to gain insight into the GTP state of microtubules. However, different analogues induce distinct microtubule structures with different stability ^8-11^. Moreover, EBs that bind autonomously to growing microtubule ends and are often considered markers of the protective cap ^12-18^, bind with varying affinities to microtubules grown in different GTP analogues ^18, 19^. Therefore, it is still unclear what the GTP microtubule conformation at growing microtubule ends looks like and whether EBs do indeed recognize the GTP state. Pure GTP microtubules have so far not been generated.

To examine the relationship between GTP hydrolysis and conformational changes in the GTP cap and to answer the question of the nucleotide preference of EBs, we produced recombinant human tubulin, which allowed us to alter the GTPase activity of microtubules. We generated wild-type (wt) α/β-tubulin and a tubulin variant with blocked GTP hydrolysis by mutating the evolutionarily conserved catalytic glutamate 254 of α-tubulin ^20^ to alanine (E254A) (Fig. 1a-b, Suppl. Fig. 1). This mutation is lethal in budding yeast, as cell division fails due to loss of microtubule dynamics ^21^.

**Fig. 1:**
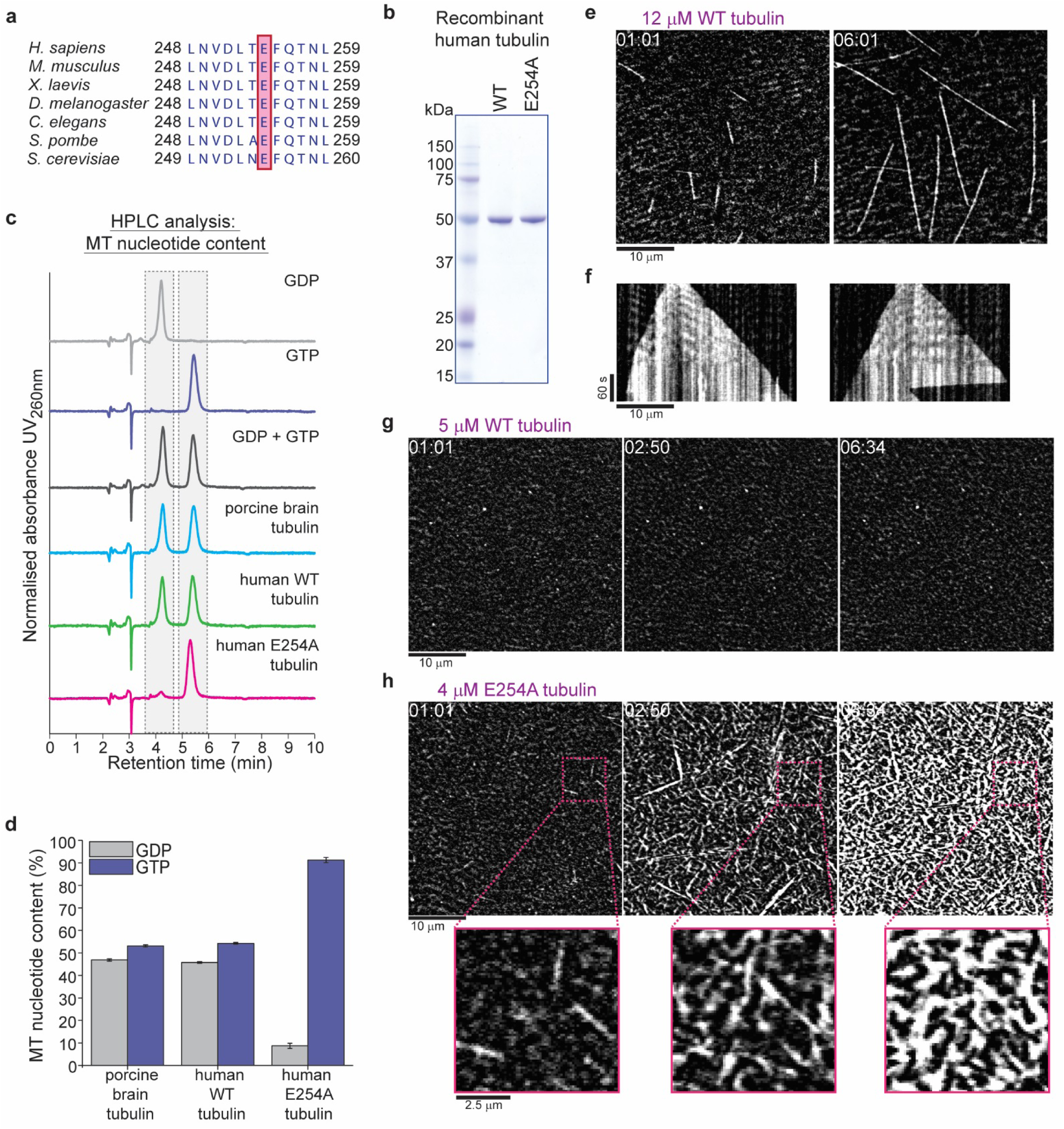
Recombinant human tubulin lacking GTPase activity strongly nucleates microtubules. **(a)** The evolutionarily conserved catalytic glutamic acid of α-tubulin (E, pink). **(b)** Coomassie Blue-stained SDS gel of purified wild type (wt) and E254A mutant recombinant human tubulin. **(c)** HPLC chromatograms of pure GDP and GTP compared to nucleotides extracted from microtubules polymerized from porcine brain tubulin or wt and E254A mutant human tubulin. **(d)** Quantification of the nucleotide content of different microtubules. Bar graphs depict the means of 3 independent experiments, error bars represent standard deviation (SD). **(e)** iSCAT microscopy images of unlabeled wt human tubulin (12 µM) growing from immobilized GMPCPP-stabilized biotinylated seeds (GMPCPP-seeds). **(f)** Kymographs showing wt microtubule growth, conditions as in (e). **(g, h)** iSCAT microscopy images showing **(g)** lack of microtubule nucleation at 5 µM wt tubulin and **(h)** strong nucleation at 4 µM E254A tubulin at a surface with an immobilized rigor kinesin (Kin1^rigor^). Bottom: magnified E254A microtubule views. Scale bars as indicated, time is always min:sec.

We measured the GTP content of microtubules polymerized from purified recombinant human wt tubulin and tubulin purified from porcine brain as a control. We observed the expected 1:1 ratio of GTP:GDP (Fig. 1c-d), since GTP is known to be hydrolyzed only at its inter-tubulin dimer, but not at the intra-dimer binding site ^22^. In contrast, E254A microtubules were almost exclusively GTP bound (Fig. 1c-d), demonstrating that GTP hydrolysis is indeed blocked in this mutant.

To visualize the growth dynamics of label-free recombinant microtubules elongating from surface-attached stable microtubule ‘seeds’, we used interferometric scattering (iSCAT) microscopy (Suppl. Fig. 2a). In the presence of GTP, human wt microtubules grew dynamically similarly to porcine brain microtubules (Fig. 1e-f, Suppl. Fig. 2b-c, Movie S1), displaying occasional transitions to depolymerization, called catastrophes, as expected ^23, 24^.

**Fig. 2:**
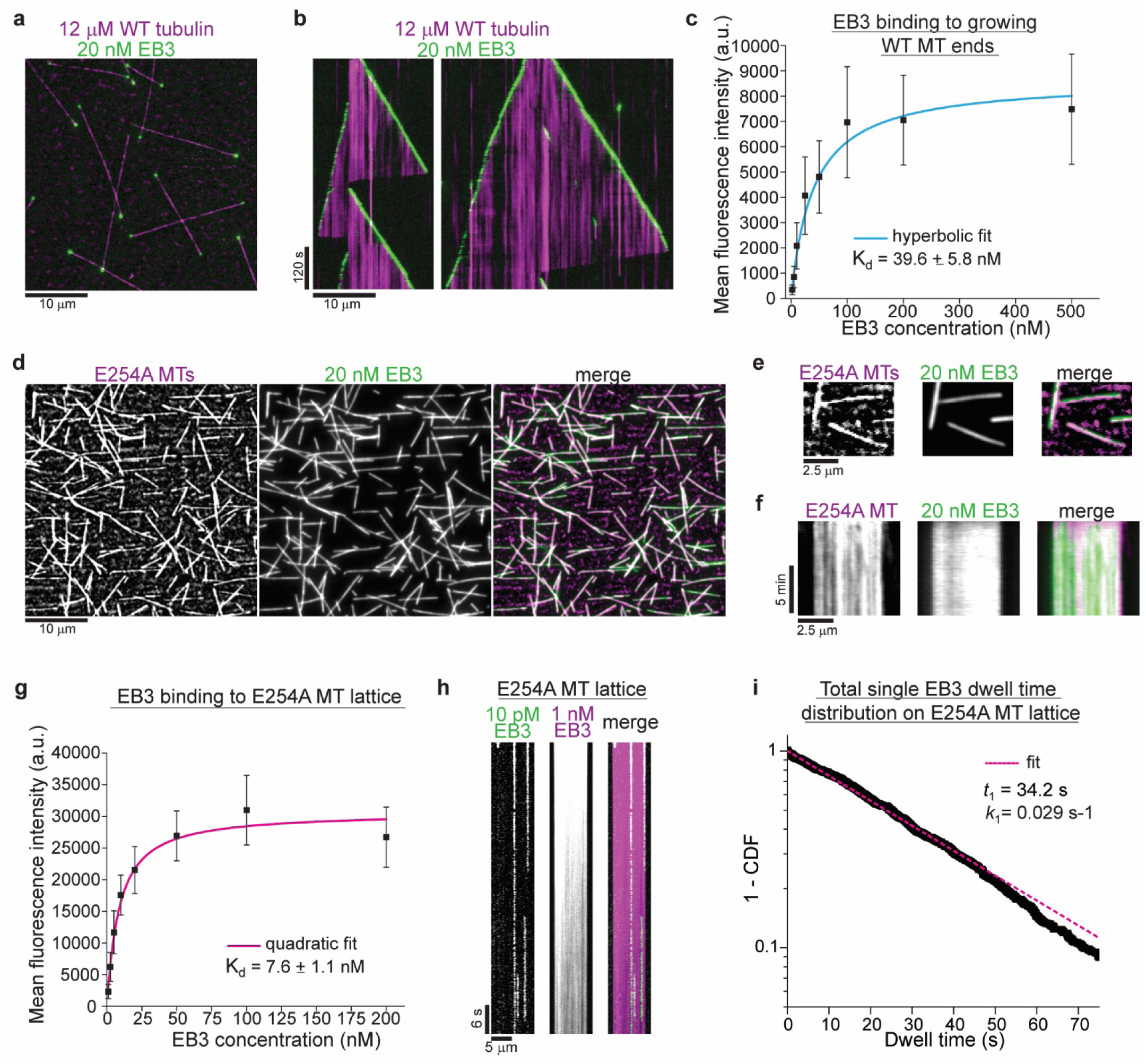
GTPase deficient microtubules are stable and bind EB3 with high affinity. **(a)** iSCAT/TIRF microscopy image of unlabeled wt microtubules (magenta) growing from GMPCPP-seeds at 12 µM wt human tubulin in the presence of 20 nM human mGFP-EB3 (green). **(b)** Kymographs showing mGFP-EB3 (green) tracking the ends of growing wt microtubules, conditions as in (a). **(c)** Quantification of mGFP-EB3 binding affinity to microtubule ends growing at 12 µM wt tubulin. Black symbols depict the averaged maximal mGFP intensities at growing plus ends at each condition, error bars represent SD. A hyperbolic fit (cyan) was used to calculate the Kd. Number of microtubules (and frames) averaged for each mGFP-EB3 concentration: 2 nM – 64 (7360), 5 nM – 87 (9222), 10 nM – 93 (8742), 25 nM – 89 (8900), 50 nM – 56 (5600), 100 nM – 58 (4756), 200 nM – 56 (4088), 500 nM – 46 (4140). **(d)** iSCAT/TIRF microscopy images of 20 nM mGFP-EB3 (green) binding to unlabeled E254A mutant microtubules (magenta) attached to a Kin1^rigor^ surface. **(e)** Magnified images and **(f)** kymographs of stable unlabeled E254A microtubules (magenta) decorated with 20 nM mGFP-EB3 (green), conditions as in (d). **(g)** Quantification of mGFP-EB3 binding affinity to E254A mutant microtubules. Black symbols depict the averaged mGFP intensities measured all along the microtubules at each condition, error bars represent SD. A quadratic fit (magenta) was used to calculate the K_d_. Number of microtubules measured for each mGFP-EB3 concentration: 1 nM – 96, 2.5 nM – 126, 5 nM – 86, 10 nM – 69, 20 nM – 78, 50 nM – 61, 100 nM – 64, 200 nM - 75. **(h)** Kymograph of mGFP-EB3 (green) single molecule imaging at 10 pM on an individual E254A microtubule in the presence of 1 nM Alexa647-EB3 (magenta). **(i)** Dwell time analysis of mGFP-EB3 molecules plotted as a survival function (1 - CDF, cumulative density function). Dashed line (magenta) is a mono-exponential fit to the data (black dots). Number of microtubules analyzed - 126, number of mGFP-EB3 binding events - 807. Scale bars as indicated.

Whereas wt tubulin hardly nucleated any microtubules spontaneously (Fig. 1e-g, Supplementary Videos 1, 2), E254A tubulin nucleated microtubules very efficiently even at low concentrations, quickly reaching high microtubule densities (Fig. 1h, Supplementary Video 3). Moreover, these mutant microtubules were hyper-stable for hours, never exhibiting catastrophes. These properties agree with the anticipated behavior of hydrolysis-deficient microtubules, and are similar to the behavior of microtubules polymerized in the presence of the non-hydrolysable GTP analogue GMPCPP ^25^.

Next, we tested how EBs bind to human wt and to hydrolysis-deficient microtubules. Simultaneous total internal reflection fluorescence (TIRF) microscopy of mGFP-EB3 (Suppl. Fig. 3a) and iSCAT imaging of unlabeled microtubules revealed that EB3 decorated growing plus and minus ends of wt microtubules in the expected comet-like manner (Fig. 2a-b, Supplementary Video 4) ^12, 13, 16^. In clear contrast, EB3 decorated the entire lattice of GTPase-deficient E254A mutant microtubules (Fig. 2d-f), reminiscent of its binding to microtubules grown in the presence of the non-hydrolysable GTP analogue GTPγS ^19^. EB3 binding to E254A microtubules was strong with an apparent dissociation constant of ∼ 8 nM (Fig. 2g) compared to ∼ 40 nM measured at growing wt ends (Fig. 2c, Suppl. Fig.3b-c). Moreover, while EB3 showed high affinity for E254A microtubules, it failed to bind human wt microtubules polymerized in the presence of the other GTP analog, GMPCPP (Suppl. Fig. 3d-e), as observed previously with mammalian brain microtubules ^18, 19^. Unbinding of mGFP-EB3 from E254A microtubules was very slow (Fig. 2h-i, Suppl. Fig.3f-g). Together these results show that EB3 displays strong affinity to microtubules locked in the GTP state, clearly establishing it as a *bona fide* GTP cap marker.

**Fig. 3:**
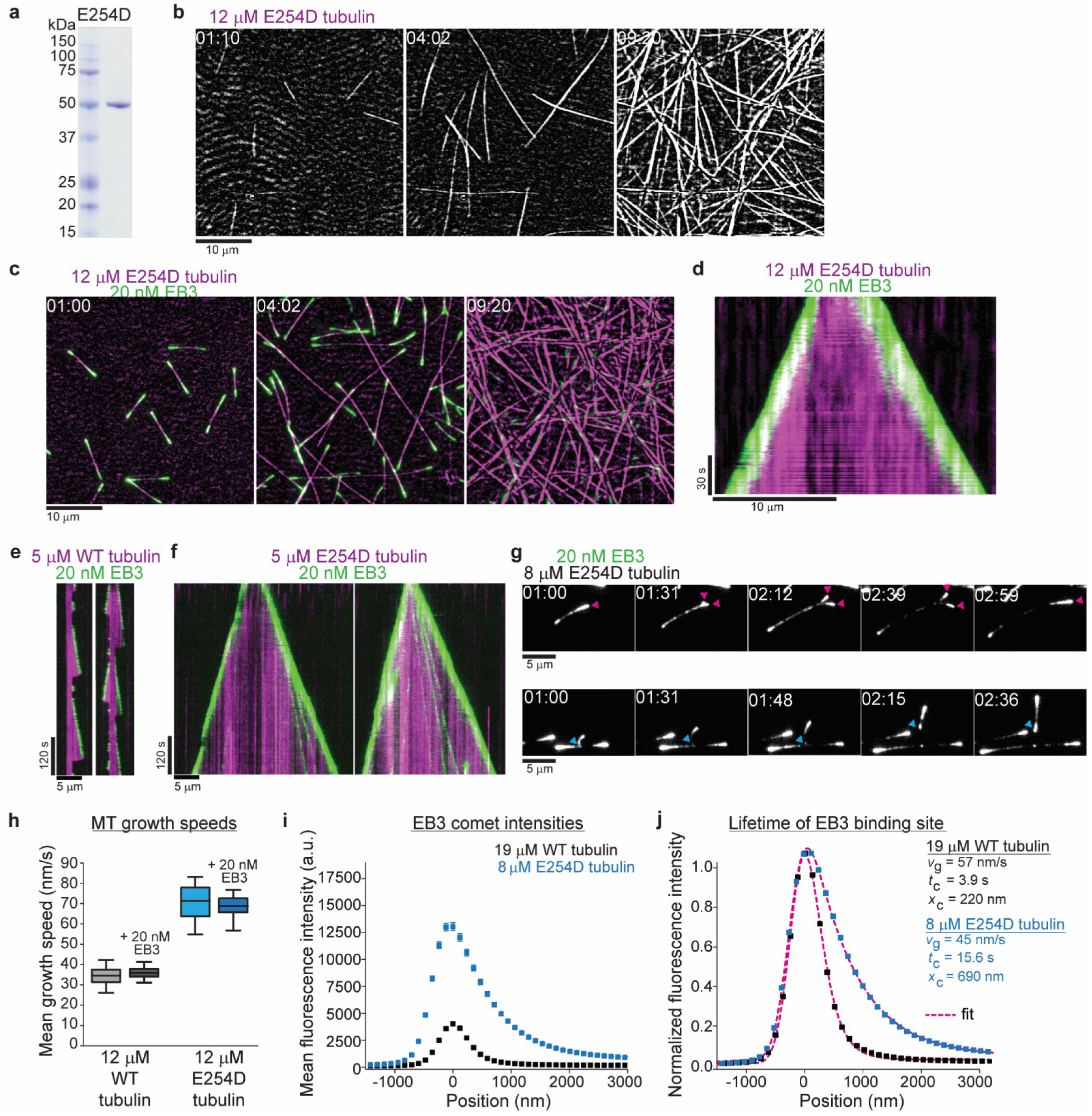
Slowing down GTP hydrolysis extends the GTP-cap and stabilizes growing microtubules. **(a)** Coomassie Blue-stained SDS gel with purified wt and E254A mutant recombinant human tubulin. **(b, c)** iSCAT/TIRF microscopy images of unlabeled E254D mutant microtubules growing from GMPCPP-seeds and also spontaneously nucleating at 12 µM E254D mutant human tubulin **(b)** in the absence and **(c)** presence of 20 nM human mGFP-EB3 (green). **(d)** Kymograph showing mGFP-EB3 (green) tracking the ends of persistently growing E254D microtubules, conditions as in (C). **(e, f)** iSCAT/TIRF microscopy kymographs depicting **(e)** wt and **(f)** E254D microtubules (magenta) growing at 5 µM tubulin in the presence of 20 nM mGFP-EB3 (green). E254D microtubules display very few catastrophes under conditions where wt human microtubules undergo frequent catastrophes. **(g)** Quantification of wt and E254D microtubule growth speeds at 12 µM respective tubulin concentrations in the presence and absence of mGFP-EB3. The boxes extend from 25^th^ to 75^th^ percentiles, the whiskers extend from 5^th^ to 95^th^ percentiles, and the mean value is plotted as a line in the middle of the box. Number of microtubules measured for each condition: 12 µM wt tubulin – 95, 12 µM wt tubulin and 20 nM mGFP-EB3 – 86, 12 µM E254D tubulin – 35, 12 µM E254D tubulin and 20 nM mGFP-EB3 - 83. **(h)** Mean mGFP-EB3 intensity profiles at growing wt (black) and E254D (blue) microtubule plus ends. Number of microtubules (and frames averaged) for each condition: wt – 68 (36960), E254D – 74 (41720). Error bars are SE. (**i)** Normalized comet profiles from (H) with dashed lines (magenta) representing exponentially modified Gaussian fits (see Methods) yielding the comet length *x*_c_ and, knowing the measured growth speed *v_g_*, the life time *t*_c_ = *x_c_*/*v_g_* of the EB binding sites. **(j)** TIRF microscopy images of protofilament bending, and re-association (arrowheads, magenta) or splitting (arrowheads, cyan) at unlabeled E254D microtubule ends growing at 8 µM tubulin visualized by mGFP-EB3. Scale bars as indicated, time is min:sec.

E254A microtubules locked in the GTP state simultaneously capture properties that different GTP analogues induce separately in wild type microtubules: strong microtubule nucleation as in the presence of GMPCPP, and strong EB binding as in the presence of GTPγS ^19, 25^. This indicates that microtubules grown in GTP analogues only display partial aspects of the GTP conformation of microtubules.

To investigate how GTP hydrolysis determines conformational transformations within the GTP cap, we generated a human tubulin mutant, in which glutamate 254 is substituted by aspartate (E254D) (Fig. 3a). This chemically similar but shorter catalytic residue is expected to slow down GTP hydrolysis compared to wt microtubules. E254D microtubules grew steadily from surface-immobilized seeds (Fig. 3b), hardly ever displaying any catastrophes even under conditions where wt microtubules displayed frequent depolymerization events (Fig. 3e-f, Supplementary Videos 5–7). E254D tubulin also nucleated microtubules more efficiently in solution than wt tubulin, but less than E254A tubulin (Fig. 3b), indicating an intermediate stability of E254D microtubules compared to wt and completely GTPase-deficient E254A microtubules. This suggests that GTP hydrolysis is indeed slowed down in the E254D mutant.

EB3 tracked growing E254D microtubule ends, displaying strikingly longer and brighter fluorescent ‘comets’ (Fig. 3c-d, Supplementary Video 5 – 7), again clearly indicative of slower GTP hydrolysis. E254D microtubules grew about twofold faster than wt, either with or without EB3 (Fig. 3g). A comparison of the average EB3 intensity profiles along the ends of wt and E254D microtubules growing at tubulin concentrations adjusted such that they grow with comparable speeds, shows that around ∼5 more EB3 binds to the ends of E254D microtubules (Fig. 3h). This increase can to a large extent be explained by the longer EB3 binding region on the mutant microtubules (Fig. 3i). Quantitative analysis of the intensity profiles allows to extract the characteristic comet lengths and, knowing the measured growth speeds, also the lifetime of the EB3 binding sites ^12, 13, 17, 26^. This revealed a four-fold increase in the lifetime of the EB3 binding sites at the ends of E254D microtubules compared to wt microtubules (Fig 3i). These results demonstrate that slowed-down GTP hydrolysis extends the lifetime and hence the size of the GTP cap.

We occasionally observed ‘split comets’ at the growing ends of the E254D microtubules that could later re-join (Fig. 3j, magenta arrowheads) or detach and grow individually (Fig. 3j, cyan arrowheads). This indicates that even partial microtubule end structures can be more stable when GTP hydrolysis is slow. This is reminiscent of recently observed split microtubule ends in the presence of the protofilament capping drug eribulin and the microtubule-stabilizing protein CLASP ^27, 28^. The ‘curved’ appearance of these partial comets likely reflects global conformational differences between unfinished end structures and fully formed tubes, pointing to a high degree of structural plasticity of the GTP cap region ^2, 29^.

How the GTP cap stabilizes growing microtubule ends is an open question. As GTP is hydrolyzed over time, a nucleotide state gradient may form at the growing microtubule end that might translate into a conformational stability gradient. Such a gradient would be detectable as a gradient of EB affinities within the GTP cap. We therefore probed the microtubule conformation along the length of the GTP cap by measuring the binding strength of EB3 at different positions within the GTP cap using single molecule imaging and dwell time analysis (Suppl. Fig. 4). We first took advantage of the elongated GTP caps of growing E254D microtubules.

We imaged single mGFP-EB3 molecules binding in the elongated cap region of growing E254D microtubules, with their end regions labeled by excess Alexa647-EB3 (Fig. 4a). We observed an overall complex (non-mono-exponential) distribution of mGFP-EB3 dwell times in the GTP cap, indicative of the presence of different conformational states with different EB3 binding affinities in the cap (Fig. 4b). Next, we performed a spatially resolved dwell time analysis by generating several ‘local dwell time distributions’ for distinct distances from the growing microtubule end. These distributions were strikingly mono-exponential displaying varying characteristic dwell times (Fig. 4e). These characteristic dwell times, and hence the EB3 binding strength, decreased gradually along the length of the GTP cap (Fig. 4e-f, Suppl. Fig. 5a), suggesting that there is a gradual conformational transition within the microtubule lattice as GTP is hydrolyzed (Suppl. Discussion). Long EB3 dwell times at the tip of the GTP cap are indicative of a more GTP-like state, given that EB3 binds strongly to GTPase deficient E254A microtubules (Fig. 2h-i), which then transforms into the lower affinity GDP state concomitant with GTP hydrolysis. The observed concerted conformational change along the length of the GTP cap most likely reflects a stability gradient whose characteristic length corresponds to the cap length (Fig. 4d).

**Fig. 4:**
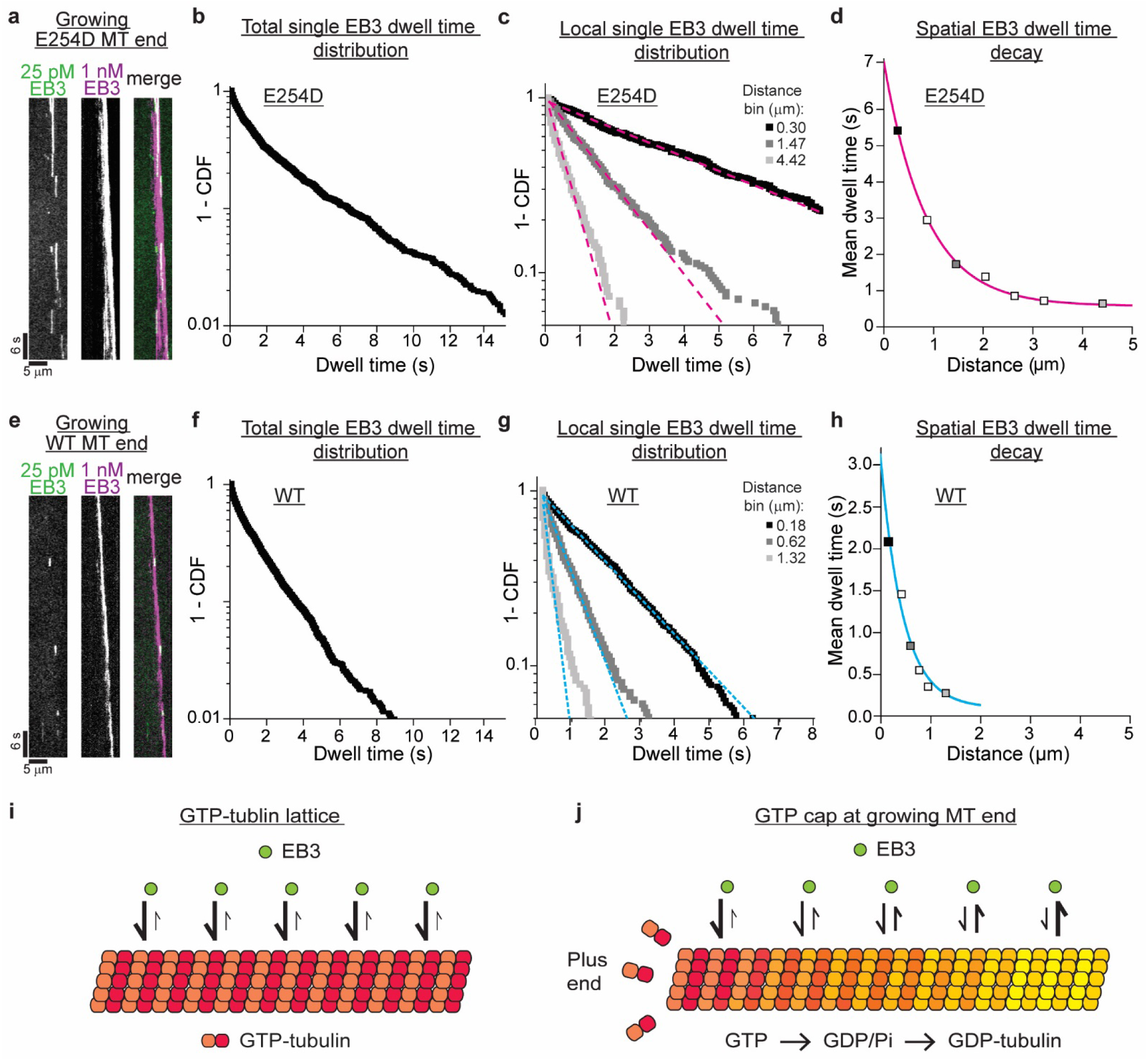
Conformational plasticity of the GTP-cap revealed by spatially resolved single EB3 dwell time distributions. **(a)** TIRF microscopy kymograph of single mGFP-EB3 molecules (green) at 25 pM in the plus end region of an E254D microtubule growing at 10 µM E254D tubulin (*v_g_* = 58.5 nm/s) in the additional presence of 1 nM Alexa647-EB3 (magenta) (for end region visualization). **(b)** Dwell time distribution of single mGFP-EB3 molecules plotted as a survival function (1-CDF, cumulative density function). Dashed line (magenta) is a mono-exponential fit to the data (black dots). Number of microtubules analyzed - 150, number of mGFP-EB3 binding events - 1834. **(c)** Local dwell time distributions at distinct distances from the growing microtubule plus end. Dashed magenta lines are mono-exponential fits. The distance bins are 0-0.59, 1.18-1.77 and 3.84-5.02 µm; bin centers shown in the legend. Source data as in (b). (**d)** Local mean dwell times as a function of distance from the growing E254D microtubule end. Filled symbols correspond to the data in (c). The solid magenta line is a mono-exponential fit. **(e)** TIRF microscopy kymograph of single mGFP-EB3 molecules (green) at 25 pM in the plus end region of a wt microtubule growing at 19 µM wt tubulin (*v_g_* = 59.8 nm/s) in the presence of additional 1 nM Alexa647-EB3 (magenta). **(f)** mGFP-EB3 dwell time distribution. Dashed line (magenta) is a mono-exponential fit to the data (black dots). Number of microtubules analyzed - 548, number of mGFP-EB3 binding events - 2425. (**g)** Local mGFP-EB3 dwell time distributions. Dashed magenta lines are mono-exponential fits. Source data as in (f). (**h)** Local mean dwell times in the wt microtubule plus end region. Filled symbols correspond to the conditions in (G). The solid magenta line is a mono-exponential fit to the data. (**i)** Schematic of high affinity EB3 binding to the GTP lattice of a GTPase-deficient microtubule. **(j)** Schematic of a growing end of a GTPase-competent microtubule displaying a gradually decreasing EB3 binding affinity as the lattice conformation gradually changes as a consequence of GTP hydrolysis.

We observed a similar gradual conformational change in the shorter GTP caps of wt microtubules growing at a similar speed (Suppl. Fig. 6), when we recorded a considerably larger number of binding events compared to previous studies ^12-14, 19, 26^ to enable spatially resolved dwell time analysis. We detected also here multiple conformational states within the GTP cap, as indicated by a complex EB3 dwell time distribution for all events observed in the entire cap region, whereas single conformational states were detected at distinct positions within the GTP cap, as indicated by roughly mono-exponential local dwell time distributions (Fig. 4f-g). The mean dwell times, indicative of the EB binding affinity, decreased over a shorter distance in wt GTP caps (Fig. 4h, Suppl. Fig.5b) than in E254D caps (Fig. 4d, Suppl. Fig. 5a), reflecting the shorter EB comet length compared to EB comets at microtubule ends with reduced GTPase activity (Fig. 3g, Suppl. Fig. 5b, 5d). This conformational gradient appears to be the response of an adaptable or ‘plastic’ microtubule lattice to GTP hydrolysis, apparently integrating a gradient of nucleotide states in the growing microtubule end region ^7, 23, 30-34^.

Taken together, our results demonstrate that the kinetics of GTP hydrolysis critically determine the properties of the GTP cap and hence microtubule stability. The ‘normal’ GTP hydrolysis rate may therefore be under evolutionary selection pressure to ensure the appropriate basic stability of microtubules in a given organism. Basic microtubule stability is linked via the GTP cap to the size of the regulatory EB binding platform that further modulates microtubule stability and interactions with target structures in the cell. Future challenges will be to elucidate the high-resolution structures of microtubules in their true GTP state and of the conformational transitions within the GTP cap to decipher the determinants of microtubule stability and function at atomic resolution.

## ACKNOWLEDGEMENTS

We thank Tarun Kapoor for the original wt tubulin expression construct, Steve Howell from the Mass Spectrometry Proteomics Science Technology Platform and the Structural Biology Science Technology Platform of the Francis Crick Institute for support, Ivo Telley and Jamie Rickman for helpful discussions, and Peter Bieling, Franck Fourniol, Nate Goehring and Sebastian Maurer for critical reading of the manuscript.

## AUTHOR CONTRIBUTIONS

J.R. and T.S. developed the study. J.R. and C.T. generated the reagents. J.R. designed and performed the experiments. The HPLC experiment was performed together with S.K. and

I.A.T. N.I.C. developed iSCAT microscopy imaging. J.R., S.K., N.I.C. and T.S. analyzed the data. J.R. and T.S. wrote and assembled the manuscript with support from N.I.C.

## DATA AND MATERIALS AVAILABILITY

All data are available in the manuscript or the supplementary materials. Correspondence regarding data and materials should be addressed to T.S.

## SUPPLEMENTARY MATERIALS AND METHODS

### Bacteria and insect cells

Bacterial strains (*Escherichia coli* DH5α, BL21 pRil, DH10MultiBac) were grown in Luria Bertani (LB) medium in the presence of appropriate antibiotics.

*Spodoptera frugiperda* strain *Sf*21 insect cells were grown in suspension at 27°C in Sf-900™ III SFM (1x) Serum Free Medium (Gibco). High Five™ cells were grown in suspension at 27°C in ExpressFive® SFM media (Gibco) supplemented with 16 mM L-glutamine (Gibco). Absence of mycoplasma in insect cell cultures was confirmed regularly.

**Molecular cloning**

To generate an expression construct for wild type human tubulin a pFastBacDual based vector containing insect cell codon-optimized versions of human α-tubulin TUBA1B (NP_006073.2) and β-tubulin TUBB3 (NP_006077.2) ^1^ (gift from T. Kapoor) was modified by Quickchange mutagenesis as follows. The fragment encoding for the deca-histidine tag and the alanine-proline linker was removed from the N-terminus of the TUBA1B. Instead, a hexa-histidine tag encoding fragment was inserted into the internal acetylation loop of the TUBA1B between isoleucine 42 and glycine 43, a strategy used previously by ^2, 3^. This resulted in the following expression construct: TUBA1B-intHis_6_ TUBB3-Gly_2_-Ser-Gly_2_-TEVsite-StrepTagII (pJR374), where the StrepTagII at the C-terminus of the TUBB3 could be removed by cleavage with Tobacco Etch Virus (TEV) protease. Removal of the N-terminal tag was necessary to improve the polymerization competency of the recombinant tubulin. The new construct was subjected to further Quickchange mutagenesis of TUBA1B to produce expression constructs for GTP hydrolysis deficient (TUBA1B^E254A^-intHis6 TUBB3-Gly_2_-Ser-Gly_2_-TEVsite-StrepTagII, pJR375) and GTP hydrolysis compromised (TUBA1B^E254D^-intHis_6_ TUBB3-Gly_2_-Ser-Gly_2_-TEVsite-StrepTagII, pJR376) versions of human tubulin dimers. These proteins are referred to as wt, E254A and E254D tubulin throughout the manuscript.

To generate a bacterial expression construct for human EB3, the coding sequence of full-length human EB3 and a long N-terminal linker were amplified by PCR from pET28a-His-mCherry-EB3 ^4^ (gift from M. Steinmetz). This fragment was fused with a monomeric GFP (mGFP) ^5, 6^ sequence in a pETMZ vector to generate a bacterial expression construct encoding for the following fusion protein: His_6_-Ztag-TEVsite-mGFP-linker-EB3 (from here on mGFP-EB3). The N-terminal His_6_-Ztag could be removed by TEV protease treatment. The expression construct for SNAP-tagged human EB3 (SNAP-EB3) has been described elsewhere ^7^. All constructs were verified by DNA sequencing.

### Protein expression

Baculovirus preparation for recombinant human tubulin expression was carried out according to manufacturer’s protocols (Bac-to-Bac system, Life Technologies) using *E. coli.* DH10MultiBac and *Sf*21 insect cells. Baculovirus-infected insect cells (BIICs) were then frozen prior to cell lysis to generate stable viral stocks as described previously ^8^.

Recombinant human tubulin expression was induced by adding 6 ml of frozen BIICs per litre to a High Five™ insect cell culture grown to densities of ∼ 1.25−10^6^ cells/ml. Cells were harvested 72 h post-induction by centrifugation (15 min, 1000 *g,* 4°C). Cell pellets were then washed with ice-cold 1x PBS, centrifuged again (15 min, 1000 *g,* 4°C), frozen in liquid nitrogen, and stored in −80°C.

mGFP-EB3 and SNAP-EB3 were expressed in *Escherichia coli* BL21 pRIL as described previously ^7^. Biotinylated monomeric *Drosophila melanogaster* kinesin-1 rigor mutant (Kin1^rigor^) was expressed in *Escherichia coli* BL21 pRIL as described previously ^9^.

### Protein purification

High Five™ insect cell pellets from 2 liters of culture expressing human recombinant tubulin were resuspended in ice-cold lysis buffer (80 mM PIPES, 1 mM EGTA, 6 mM MgCl_2_, 50 mM imidazole, 100 mM KCl, 2 mM GTP, 1 mM 2-mercaptoethanol (2-ME), pH 7.2) supplemented with protease inhibitors (Roche), DNase I (10 µg ml/ml, Sigma), using the same volume of buffer as the cell pellet. Resuspended cells were lysed by dounce homogenization (60 strokes). The lysate was then diluted 4-fold with dilution buffer (80 mM PIPES, 1 mM EGTA, 6 mM MgCl_2_, 50 mM imidazole, 2 mM GTP, 1 mM 2-ME, pH 7.2) and clarified by ultracentrifugation (158,420x *g,* 1 h, 4°C). The supernatant was passed through a 5 ml HisTrap HP column (GE Healthcare). The column was first washed with 5 column volumes (CVs) of lysis buffer, then with 5 CVs of Ni wash buffer 1 (80 mM PIPES, 1 mM EGTA, 11 mM MgCl_2_, 2 mM GTP, 5 mM ATP, 1 mM 2-ME, pH 7.2), then with 5 CVs of Ni wash buffer 2 (80 mM PIPES, 1 mM EGTA, 5 mM MgCl_2_, 0.1% (vol/vol) Tween-20, 10% (w/vol) glycerol), 2 mM GTP, 1 mM 2-ME, pH 7.2), and then again with 5 CVs of lysis buffer. The protein was eluted with Ni elution buffer (80 mM PIPES, 1 mM EGTA, 5 mM MgCl_2_, 500 mM imidazole 2 mM GTP, 1 mM 2-ME, pH 7.2). The eluate was immediately diluted 6-fold with Strep binding buffer (80 mM PIPES, 1 mM EGTA, 5 mM MgCl_2_, 2 mM GTP, 1 mM 2-ME, pH 7.2) and passed through serially connected 1 ml HiPrep SP FF and 5 ml StrepTag HP columns (both GE Healthcare) equilibrated in Strep binding buffer. The columns were then washed with 2 CVs of Strep binding buffer. The protein was eluted in Strep elution buffer (80 mM PIPES, 1 mM EGTA, 4 mM MgCl_2_, 2.5 mM D-desthiobiotin, 50 mM imidazole, 2 mM GTP, 1 mM 2-ME, pH 7.2). The eluate was diluted 2-fold in Strep elution buffer, transferred on ice and incubated with TEV protease for 2 h to remove the C-terminal StrepTagII from TUBB3. Following TEV cleavage the eluate was centrifuged at 204,428 *g,* 10 min, 4°C. The supernatant was then passed through a further 1 ml HiPrep SP FF column equilibrated in SP wash buffer (80 mM PIPES, 1 mM EGTA, 5 mM MgCl_2_, 2 mM GTP, pH 6.8) to remove TEV protease. Finally, the flow through now containing the purified recombinant human tubulin was passed through two serially connected pre-equilibrated HiPrep Desalting columns (GE Healthcare) to exchange the buffer to tubulin storage buffer (80 mM PIPES, 1 mM EGTA 1 mM MgCl_2_, 0.2 mM GTP, pH 6.8). The tubulin containing fractions were pooled, concentrated (Vivaspin 30,000 MWCO, Sartorius) to above 3.5 mg/ml, ultracentrifuged (278,088 *g*, 10 min, 4°C), aliquoted, snap frozen, and stored in liquid nitrogen until use. Mass spectroscopy demonstrated that the purified human tubulin was free of insect cell tubulin (Suppl. Fig. 1d).

mGFP-EB3 and SNAP-EB3 were purified and SNAP-EB3 was labeled with SNAP-Surface-AlexaFluor647 (NEB) as described (referred to as Alexa647-EB3 throughout the manuscript) ^7^. Biotinylated monomeric *Drosophila melanogaster* kinesin-1 rigor mutant (Kin1^rigor^) was purified as described previously ^9^.

Porcine brain tubulin was purified following a published protocol ^10^. Purified porcine brain tubulin was recycled and labeled with Alexa647-*N*-hydroxysuccinimide ester (NHS; Sigma-Aldrich), CF640R-NHS (Sigma-Aldrich), Atto565-NHS (Sigma-Alrich) or biotin-NHS (Thermo Scientific), as described previously ^11^.

EB3 concentration was determined by Bradford assay, values refer to monomer concentration. Tubulin concentration was determined by UV/VIS spectroscopy measurements (absorption at 280 nm). Values refer to tubulin dimer concentration.

### Determination of microtubule nucleotide content

To determine their nucleotide content, microtubules were polymerized from different types of tubulin in the presence of a low concentration of short microtubule ‘seeds’ to initiate microtubule growth.

First microtubule ‘seeds’ were polymerized from 15 µM recycled porcine brain tubulin in the presence of 0.5 mM GMPCPP (Jena Bioscience) in BRB80 (80 mM PIPES, 1 mM EGTA, 1 mM MgCl_2_, pH 6.8) at 37°C for 50 min. The seeds were centrifuged at 17,000 *g* at room temperature for 10 min, washed in warm BRB80, centrifuged at 17,000 g at room temperature, and resuspended in BRB80 to an estimated concentration of 15 µM tubulin assuming 100% nucleation and polymerization efficiency.

160 µg of purified porcine brain tubulin, human wt tubulin, human E254A tubulin was then used for each polymerization reaction. The polymerization reaction consisted of 20 µM tubulin diluted in BRB80 and mixed with 26% glycerol (vol/vol) and 1 mM GTP (final concentrations). These reactions were first incubated on ice for 5 min, then transferred to 37°C, and after 1 min supplemented with 0.15 µM GMPCPP-stabilized ‘seeds’, and polymerized for 50 min. The samples were then centrifuged at 278,088 x *g* for 10 min at 37°C. The supernatants were aspirated, the pellets containing microtubules washed twice with 2x sample volume of warm BRB80, and resuspended in 10 mM Tris-HCl (pH 7.5). Nucleotides were then extracted by addition of 0.7% (w/vol) HClO_4_ followed by immediate neutralization of the reaction by addition of Na-acetate to 200 mM (final concentrations). The supernatants containing the extracted nucleotides were separated from precipitated protein by centrifugation at 15,000 *g* for 5 minutes at 4°C and subsequently filtered through a 0.22 µm centrifugal filter (Durapore-PVDF, Millipore).

The samples (55 µl) were applied to a Zorbax SB-C18 column (4.6 x 250 mm, 5 µm pore size, Agilent Technologies) mounted on a Jasco HPLC system controlled by Chromnav software (v1.19, Jasco). The column temperature was maintained at 30°C. Nucleotides were separated under isocratic conditions in 100 mM K_2_HPO_4_/KH_2_PO_4_ (pH 6.5), 10 mM tetrabutylammonium bromide, 7% (vol/vol) acetonitrile. The absorbance was monitored using a Jasco MD-2010 Plus multi-wavelength detector. The GDP and GTP peaks were identified from comparison to the retention times of pure nucleotide standards (Fig. 1c). The relative amounts of GDP and GTP were determined by integrating the peaks in the 260 nm absorbance channel using Chromnav software (Fig. 1d).

### Mass spectrometry

Protein molecular mass was determined using a microTOFQ electrospray mass spectrometer (Bruker Daltonics) (Fig. 1d). Proteins were first desalted using a 2 mm x 10 mm guard column (Upchurch Scientific) packed with Poros R2 resin (Perseptive Biosystems). Protein was injected via a syringe onto the column in 10% acetonitrile, 0.10% acetic acid, washed with the same solvent and eluted in 60% acetonitrile, 0.1% acetic acid. Desalted protein was then infused into the mass spectrometer at 3 µl/min using an electrospray voltage of 4.5kV. Mass spectra were deconvolved using maximum entropy software (Bruker Daltonics).

### In vitro assays

Flow chambers were assembled similarly for all microscopy assays from poly-(L-lysine)-polyethylene glycol (PEG) (SuSoS) treated counter glass and passivated biotin-PEG-functionalized coverslips as described previously ^12^.

#### Microtubule dynamics assay

GMPCPP-stabilized biotinylated non-fluorescent microtubule ‘seeds’ (for iSCAT microscopy) or fluorescent seeds (containing 12% of CF640R-, or Atto565-labeled tubulin; for TIRF microscopy) were prepared as described previously ^9, 12^. Microtubule dynamic assays were performed as detailed earlier ^9, 12^. The passivated flow chambers were incubated for 5 min with 5% Pluronic F-127 (Sigma-Aldrich) in MQ water at room temperature, washed with assay buffer (80 mM PIPES, 1 mM EGTA, 1 mM MgCl_2_, 30 mM KCl, 1 mM GTP, 5 mM 2-ME, 0.15% (w/vol) methylcellulose (4,000 cP, Sigma-Aldrich), 1% (w/vol) glucose, pH 6.8), then with assay buffer containing κ-casein (50 μg/ml, Sigma-Aldrich). Flow chambers were then incubated on a metal block on ice in assay buffer supplemented with NeutrAvidin (50 μg/ml, Life Technologies) to coat the functionalized glass surface and washed again with assay buffer. Microtubule seeds were diluted in assay buffer, flowed into the chamber and incubated for 3 min at room temperature to allow them to attach to the NeutrAvidin-coated functionalized glass surface. The chamber was then washed again with assay buffer followed by flowing in the final assay mix. The final assay mix for the microtubule dynamics assays consisted of 98% of assay buffer containing unlabeled porcine brain, wt or mutant recombinant human tubulin (concentrations as indicated: 5 – 19 µM for different tubulins) and oxygen scavengers (180 µg/ml catalase (Sigma-Aldrich), 752 µg/ml glucose oxidase (Serva)). The final assay mix also contained 2% EB3 storage buffer (50 mM Na-phosphate, 400 mM KCl, 5 mM MgCl_2_, 0.5 mM 2-ME, pH 7.2) optionally supplemented with mGFP-EB3 (concentrations as indicated: 0 – 500 nM). The flow chamber was then sealed with silicone grease. Imaging was started 1 min after transferring the sample to the microscope chamber at 30°C.

#### Microtubule nucleation assay

Microtubule nucleation assays (Fig. 2d) were performed as described previously ^9^. The initial experimental steps to treat the flow cell were the same as in the microtubule dynamics assay. However, instead of NeutrAvidin the surface of the flow chamber was coated with biotinylated *Drosophila melanogaster* Kin1^rigor^ mutant (167 nM) by incubation in the assay buffer on a metal block on ice for 10 min. No microtubule ‘seeds’ were used. Following washes with the assay buffer, the final assay mix comprising 98% of assay buffer containing unlabeled wt or E254A mutant recombinant tubulin (concentrations indicated) and oxygen scavengers (180 µg/ml catalase, 752 µg/ml glucose oxidase) and 2% of EB3 storage buffer was flowed into the chamber.

#### Experiments with GTP-bound E254A microtubules and GMPCPP-bound wt microtubules

For preparation of stable microtubules (Suppl. Fig. 3d-e), the buffer of the wt tubulin was first exchanged from tubulin storage buffer to BRB80. Microtubules were then polymerized from 12.5 µM wt tubulin in the presence of 0.5 mM GMPCPP in BRB80 at 37°C for 1 h. The GMPCPP-stabilized wt microtubules were centrifuged at 17,000 *g* at room temperature for 10 min, washed in warm BRB80, centrifuged at 17,000 g at room temperature, and resuspended in BRB80. E254A microtubules were polymerized under the same conditions except in the presence of GTP (1 mM) instead of GMPCPP. These GTP-bound E254A microtubules remained stable throughout the day.

The initial sample preparation for imaging and flow cell treatment were identical to the nucleation assay, except that now either stable GMPCPP-stabilized wt or GTP-bound E254A microtubules were attached to the Kin1^rigor^-surface by 3 min incubation at room temperature. The chamber was then washed with assay buffer to remove unbound microtubules followed by flowing in the final assay mix consisting of 98% of assay buffer with oxygen scavengers (180 µg/ml catalase, 752 µg/ml glucose oxidase) and 2% EB3 storage buffer optionally supplemented with 20 nM mGFP-EB3.

To determine the binding affinity of mGFP-EB3 for E254A microtubules (Fig. 2g), E254A microtubules were first polymerized at 1 µM E254 tubulin from 0.5 µM GMPCPP-stabilized biotinylated fluorescently labeled porcine brain microtubule ‘seeds’ in BRB80 for 1 at 37°C for 1 h. These microtubules were then transferred to room temperature and stored until use. The initial sample preparation steps were identical to the dynamic microtubule assay, except that now the GMPCPP-stabilized ‘seeds’ with stable unlabeled E254A-lattice extensions diluted in BRB80 were attached to the NeutrAvidin-coated functionalized glass surface. The final assay mix in these experiments consisted of 98% of assay buffer containing oxygen scavengers (180 µg/ml catalase, 752 µg/ml glucose oxidase) and 2% EB3 storage buffer supplemented with mGFP-EB3 (concentrations as indicated: 1 – 200 nM).

Samples for EB3 washout experiments with E254A microtubules (Suppl. Fig. 3f) were prepared identically to the samples for mGFP-EB3 binding affinity determination. The mGFP-EB3 concentration was 2.5 nM. After initial imaging the sample was removed from the microscope objective and washed with ∼ 20 flow chamber volumes of assay buffer. Then assay buffer containing oxygen scavengers but no mGFP-EB3 was flowed in the chamber, and the sample was imaged again to visualize the remaining mGFP-EB3 binding.

#### Single molecule assays

Single molecule experiments on dynamic microtubules were performed similarly to the dynamic assays (Fig. 4). We chose wt and E254D tubulin concentration, at which wt and E254D microtubules grew at nearly identical growth speeds (Suppl. Fig. 6). The final assay mix consisted of 98% of assay buffer containing wt tubulin (19 µM) or E254D tubulin (10 µM) and oxygen scavengers (180 µg/ml catalase, 752 µg/ml glucose oxidase) and 2% of EB3 storage buffer supplemented with 25 pM mGFP-EB3 and 1 nM Alexa647-EB3.

Single molecule experiments with E254A microtubules were carried out under identical experimental and imaging conditions as the single molecule experiments with wt and E254D microtubules, except using E254A microtubules grown from GMPCPP-stabilized ‘seeds’ (described above) instead of dynamic microtubules. The final assay mix did not include soluble tubulin but only mGFP-EB3 (10 pM) and Alexa647-EB3 (1 nM).

#### Steady state bleaching assay

Samples for steady state bleaching assays were prepared identically to the single molecule experiments for each respective tubulin species (wt, E254D, E254A). Except that only 2.5 nM mGFP-EB3 was included in the final assay mix (and no Alexa647-EB3).

### Microscopy

#### Simultaneous iSCAT and TIRF microscopy

A total internal reflection fluorescence (TIRF) microscope system (Cairn Research, UK), was modified to allow simultaneous TIRF and interferometric scattering (iSCAT) microscopy ^13^ for the detection of GFP-EB3 and unlabeled microtubules, respectively. The collimated beams from a 488 nm and 561 nm diode laser were focused on the back focal plane of a 60x 1.49 N.A. TIRF objective (Nikon, Japan): the 488 nm beam was positioned to give TIRF illumination and the 561 beam was positioned at ∼18° to give epi illumination. Both beams were rapidly scanned azimuthally using a galvo scanning system (iLas2, Gataca Systems, France) to provide uniform illumination and reduce interference effects. A quadband dichroic mirror (Chroma: ZT405/488/561/640) and band-pass filters (Chroma: ET525/50, ZET561/10) were used to separate detected fluorescence and scattered 561 nm laser light. Both channels were recorded simultaneously using two EMCCD cameras (Ixon Ultra 888, Andor, UK). Both camera relays had a total magnification of 2x. Time-lapse movies were acquired with 63 ms exposure time at 1 Hz for 10 minutes.

For each sample, an average static background image was created by translating the sample rapidly while acquiring a movie stream, and averaging the resulting image series. The raw iSCAT movie was then divided by the corresponding average background image to produce a pseudo-flat-field corrected, normalized movie. This was then filtered using a mask in Fourier-space, to remove large-scale dynamic interference effects, and a Kalman stack filter to reduce noise.

#### TIRF microscopy

For ‘TIRF microscopy only’ experiments (Fig. 2c, g-i, 3h-i, 4, Suppl. Fig.-s 3b-c, 3f-g, 5, 6) movies were acquired using a custom TIRF microscope (iMIC, FEI Munich) at 30 °C, described in detail previously ^9, 14^. Exposure times were between 55 ms (single molecule imaging) and 150 ms (nucleation assays). Images were acquired at intervals of every 60 ms (single-molecule imaging), 1 s (nucleation assays), or 200 ms (comet analysis) keeping imaging conditions consistent for each set of experiments. For steady-state bleaching, movies were acquired with identical laser illumination powers and exposure times (55 ms), and increasing frame intervals (60, 100, 200 ms).

### Data analysis

#### Image analysis

Images were processed and analyzed using the Fiji package of ImageJ (https://fiji.sc), using custom macros. Further analysis and data processing was carried out in MATLAB (MathWorks USA) and Origin (OriginLab, USA).

Kymographs were generated as described previously ^7, 15^. After acquisition, movies were corrected for drift when necessary, then individual microtubule growth trajectories were drawn on a maximum intensity projection of the entire movie stack (Suppl. Fig. 4a). Kymographs were generated from the movie stack along each trajectory line; subsequently, the microtubule end position was traced manually on each kymograph (Suppl. Fig. 4b) (*3*). Periods including overlapping microtubules were excluded from the analysis. Microtubule growth speeds were calculated from the marked end position on the kymographs.

Average spatial mGFP-EB3 intensity profiles (comets) (Fig. 3h, 3i) were generated from kymographs as described previously ^7, 15^. Each kymograph was straightened and re-centered using the marked microtubule end positions, then aligned and averaged with other straightened kymographs. The intensities were averaged for all time points at each position along the resulting average kymograph, giving a time-averaged spatial intensity profile. Kinetic rate constants were extracted from an exponentially modified Gaussian fit to the comet profile, as described previously ^14^.

Averaged maximal comet intensities were used to determine the EB3 affinity to wt microtubules (Fig. 2c). Averaged lattice binding affinities were used to determine EB3 binding affinity to E254A microtubules (Fig. 2g). Equilibrium dissociation constants (K_d_) for mGFP-EB3 binding to E254A microtubules and to wt microtubule ends were determined from a quadratic fit to to the E254A microtubule data (Fig. 2g), using a measured total concentration of 2 nM of binding sites along the E254E microtubules in these experiments (data not shown), and from a hyperbolic fit to the wt data (Fig. 2c), since in this case EB3 was in large excess over binding sites at microtubule ends.

#### Single molecule dwell time analysis

To obtain dwell time distributions for the single molecule mGFP-EB3 experiments (Figs. 2h, 2i, 4a-h), kymographs were generated and the microtubule end positions traced, as described above (Suppl. Fig. 4a-b). Using the traced end position, the corresponding x-y coordinates of the microtubule end in the original movie were calculated for each frame (Suppl. Fig. 4c). These coordinates were used to create a binary mask movie that only included points on each microtubule up to the growing end in each frame. The raw image data in each frame of a movie were analyzed with a Single Molecule Localization procedure (GDSC SMLM plugin for ImageJ: https://github.com/aherbert/GDSC-SMLM); this determined the coordinates of each potential single molecule with sub-resolution precision (typically ∼< 30nm). The resulting localization positions were filtered in space and time using the binary mask movie, to remove all events not localized on a microtubule (Suppl. Fig. 4d). The remaining localization events were linked together in space and time throughout the whole movie using specific thresholds: only two events that occurred within a maximum separation of one pixel (120 nm) and 30 frames (1.8 s) were linked together. This allowed for slight movement of the molecules between frames (due to e.g. lattice diffusion, microtubule wiggling, drift) and dark frames (due to e.g. blinking). The linking parameter values were chosen after a careful inspection of the effects of the parameter space on the resulting individual dwell events, for a specific kymograph. The automatically identified binding events agreed well with visually identified events in test subsets of the data (Suppl. Fig. 4e). For each molecule, the dwell time was determined by the total time over which events were linked. The position of the molecule in the first frame of the binding event relative to the nearest microtubule end was calculated, creating a list of dwell times with their corresponding distances from the growing microtubule end (Suppl. Fig. 4f). For spatially resolved dwell time distributions, binding events were binned at specific distances from the end (Suppl.. Fig. 4g) and the dwell time ‘1 - cumulative distribution function’ (survival function) calculated for each bin. Characteristic dwell times were extracted from these distributions using a mono-exponential fit.

#### Steady state bleaching analysis

EB3 single molecule dwell times on wild-type and E254D microtubule ends (Fig. 4) were essentially unaffected by bleaching, because they were much shorter than the bleaching time of ∼45 s (Suppl. Fig. 3g). However, single molecule dwell times on E254A microtubules were in the range of our bleaching time and were therefore determined from a steady state bleaching analysis performed at different time intervals (Suppl. Fig. 3g) ^16^.

We assume that molecules bound to the microtubule bleach at a rate *k*_b_ and unbind at a rate *k*_off_. We assume an excess of unbleached EB3 molecules in solution such that at steady-state the binding rate of unbleached molecules is limited by the total unbinding rate. Solving the resulting differential equations gives a relative unbleached fraction at time *t* after illumination of

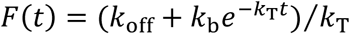

Where *k*_T_ = *k*_b_ + *k*_off_

Movies taken at different frame intervals were background corrected using a 50 pixel rolling ball subtraction. The mean intensity over the field of view was calculated for each frame, and normalized to the value in the first frame. Decay curves were fit with the function above, with *k*_off_ shared globally (Suppl. Fig. 3g).

## SUPPL. DISCUSSION

### EB3 comets

The shape of averaged mGFP-EB3 intensity profiles (EB comets) at the plus-ends of microtubules polymerizing from human wild-type or E254D tubulin can be explained by a mono-exponential decay of the EB3 density along the microtubule (Fig. 3g). For human wild-type microtubules, the characteristic comet length was in the range measured previously with porcine brain microtubules (considering similar growth speeds) ^17-23^. This confirms that microtubule lattice-incorporated recombinant human wild-type tubulin, and porcine brain tubulin, hydrolyze GTP with similar speeds ^23^. EB comets at the ends of E254D microtubules were longer than wild-type comets, in agreement with slowed down GTP hydrolysis (Fig. 3g).

Here, we did not try to determine with high precision how close the EB3 peak was to the growing microtubule end position ^14^. Our iSCAT microscopy setup allowed us to image microtubules consisting entirely of unlabeled recombinant tubulin, but did not produce images of the quality required for automated microtubule end tracking with sub-pixel precision ^24, 25^. Therefore, we based our comet and single molecule analysis entirely on TIRF microscopy movies of fluorescently labeled EB3. The kymograph-based method used here has a spatial resolution of 1-2 pixels, i.e. ∼200 nm, which was sufficient for our analysis here.

Previous comet analysis revealed a short region at the very end of the growing microtubule to which EBs did not bind ^14^. This short region could be detected using high precision tracking of microtubule ends (with sub-pixel precision) when microtubules grew very fast (faster than the highest speeds studied here). We showed previously that an EB-free zone at the extreme microtubule end could be in part explained by the EB association kinetics. We furthermore discussed that the EB-free zone could be a consequence of the particular structure of growing microtubule ends ^14^. It is now likely that EB binding sites may be absent at the very end of growing microtubules, because cryo-EM tomography has revealed in cells and in vitro the existence of individual curved protofilaments at the very ends of growing microtubules. These individual protofilaments did not form lateral interactions, and hence, did not form the binding site for EBs ^26, 27^.

Previously, we discussed the possibility that EBs might not bind well to GTP tubulins at the extreme microtubule end ^14^, because of the results obtained with GTP analogues. EBs bind well to microtubules grown in the presence of GTPγS, which has sometimes been considered to be a transition state mimic, but bind only weakly to microtubules grown in the presence of GMPCPP ^20, 21, 27^, which has typically been considered to be the best GTP mimic for a microtubule ^28-30^. However, our results with the GTPase-deficient mutant E254A tubulin clearly show here that EBs bind very well to microtubules in a true GTP state. It is likely that microtubules grown in nucleotide analogues only partially mimic the true GTP conformation, and different analogues mimic different aspects of it.

### Single molecule dwell times on wild-type microtubules

The average dwell time measured for single mGFP-EB3 molecules bound to the growing ends of recombinant human wild-type microtubules was 880 ms (Fig. 4f), which is comparable to previous reports of EB dwell times being in the range of 50-300 ms ^4, 17, 18, 20^. The slightly longer mean dwell time observed here is probably a consequence of the lower ionic strength buffer used. However, this longer dwell time helped us to unambiguously identify a spatial affinity gradient for mGFP-EB3 binding not only within the elongated GTP cap of the E254D microtubules (Fig. 4c, 4d), but also in the considerably shorter caps of wt microtubules (Fig. 4g, 4h).

### Single molecule dwell times on hydrolysis-deficient E254A microtubules

We found that mGFP-EB3 bound strongly to hydrolysis-deficient E254A microtubules. Washout experiments showed that a considerable amount of EB3 was still bound to E254A microtubules minutes after mGFP-EB3 was washed out of the flow chamber (Suppl. Fig. 3f), indicative of a dwell time in the minute range. This means that photo-bleaching likely prematurely ends a considerable fraction of the observed individual mGFP-EB3 binding events to E254A microtubules at our single molecule imaging conditions (Fig. 2h, 2i). Therefore, we imaged mGFP-EB3 at higher concentrations (above single molecule imaging conditions) in binding equilibrium with E254A microtubules over time at different time lapse intervals (Suppl. Fig. 3g). Both the bleaching and the unbinding rate could be determined from these steady-state bleaching time courses (Methods). We found an EB3 unbinding rate of 0.012 s^-1^ (0.72 min^-1^) and a bleaching rate of 0.022 s^-1^ (1.3 min^-1^) for our single molecule imaging conditions: these rates are consistent with the measured apparent mean dwell time of 0.029 s^-1^ for single GFP-EB3 molecules on E254A microtubules (Fig. 2i), which is expected to be the sum of the unbinding and the bleaching rate. In conclusion, mGFP-EB3 molecules remain bound to E254A microtubules for about a minute.

The difference between the single molecule dwell times of EB3 bound to E254A microtubules (Fig. 2i) versus wild-type microtubule ends (Fig. 4b) was considerably larger than the difference between the corresponding dissociation constants (Fig. 2c, 2g). This indicates that not only the unbinding rate, but also the binding rate is reduced on E254A microtubules compared to wild-type microtubules, even if to a lesser extent. This suggests that EB3 may induce and stabilize a conformational lattice transformation in E254A microtubules when it binds, akin to an ‘induced fit’ scenario. Such an effect on the microtubule lattice would agree with previous observations showing that EB binding to microtubules can change their protofilament twist ^27, 31, 32^ and can mildly accelerate cap maturation ^20^, which means ‘accelerate GTP hydrolysis’, in agreement with our results here as well as recent structural observations ^27^.

## SUPPLEMENTARY VIDEO LEGENDS

**Supplementary Video 1. Pure recombinant human wild-type tubulin polymerizes into dynamic microtubules.** iSCAT microscopy time-lapse movie of unlabeled microtubules growing in the presence of 12 µM human wt tubulin from immobilized stable microtubule seeds at 30°C. Scale bar is 5 µm. Time stamp is in min:s.

**Supplementary Video 2. Human wild-type tubulin does not nucleate microtubules at low tubulin concentration.** iSCAT microscopy time-lapse movie showing lack of microtubule nucleation at 5 µM human wt tubulin on a surface with an immobilized rigor kinesin (Kin1^rigor^) at 30°C. Scale bar is 5 µm. Time stamp is in min:s.

**Supplementary Video 3. The human E245A tubulin mutant lacking GTPase activity strongly nucleates microtubules.** iSCAT microscopy time-lapse movie showing strong microtubule nucleation at 4 µM human E254A tubulin on a surface with an immobilized rigor kinesin (Kin1^rigor^) at 30°C. Scale bar is 5 µm. Time stamp is in min:s.

**Supplementary Video 4. EB3 binds to the growing ends of human wild-type microtubules.** iSCAT/TIRF microscopy time-lapse movie of unlabeled wt microtubules (magenta) growing from immobilized stable microtubule seeds in the presence of 12 µM human wt tubulin and 20 nM human mGFP-EB3 (green) at 30°C. Scale bar is 5 µm. Time stamp is in min:s.

**Supplementary Video 5. Slowing down GTP hydrolysis extends the GTP-cap and stabilizes growing microtubules.** iSCAT/TIRF microscopy time-lapse movie of unlabeled E254D mutant microtubules (magenta) growing from immobilized microtubule seeds and also spontaneously nucleating in the presence of 12 µM human E254D mutant tubulin and 20 nM human mGFP-EB3 (green) at 30°C. Scale bar is 5 µm. Time stamp is in min:s.

**Supplementary Video 6. Wild-type microtubules are very dynamic at low tubulin concentrations.** iSCAT/TIRF microscopy time-lapse movie of human wt microtubules (magenta) undergoing frequent catastrophes when growing from immobilized microtubule seeds in the presence of 5 µM human wt tubulin and 20 nM human mGFP-EB3 (green) at 30°C. Scale bar is 5 µm. Time stamp is in min:s.

**Supplementary Video 7. Microtubules with reduced GTPase activity are more stable than wild-type microtubules.** iSCAT/TIRF microscopy time-lapse movie of human E254D microtubules (magenta) show persistent growth when growing from immobilized microtubule seeds in the presence of 5 µM E254D tubulin and 20 nM human mGFP-EB3 (green)at 30°C. Scale bar is 5 µm. Time stamp is in min:s.

## SUPPLEMENTARY FIGURES

**Supplementary Fig. 1:**
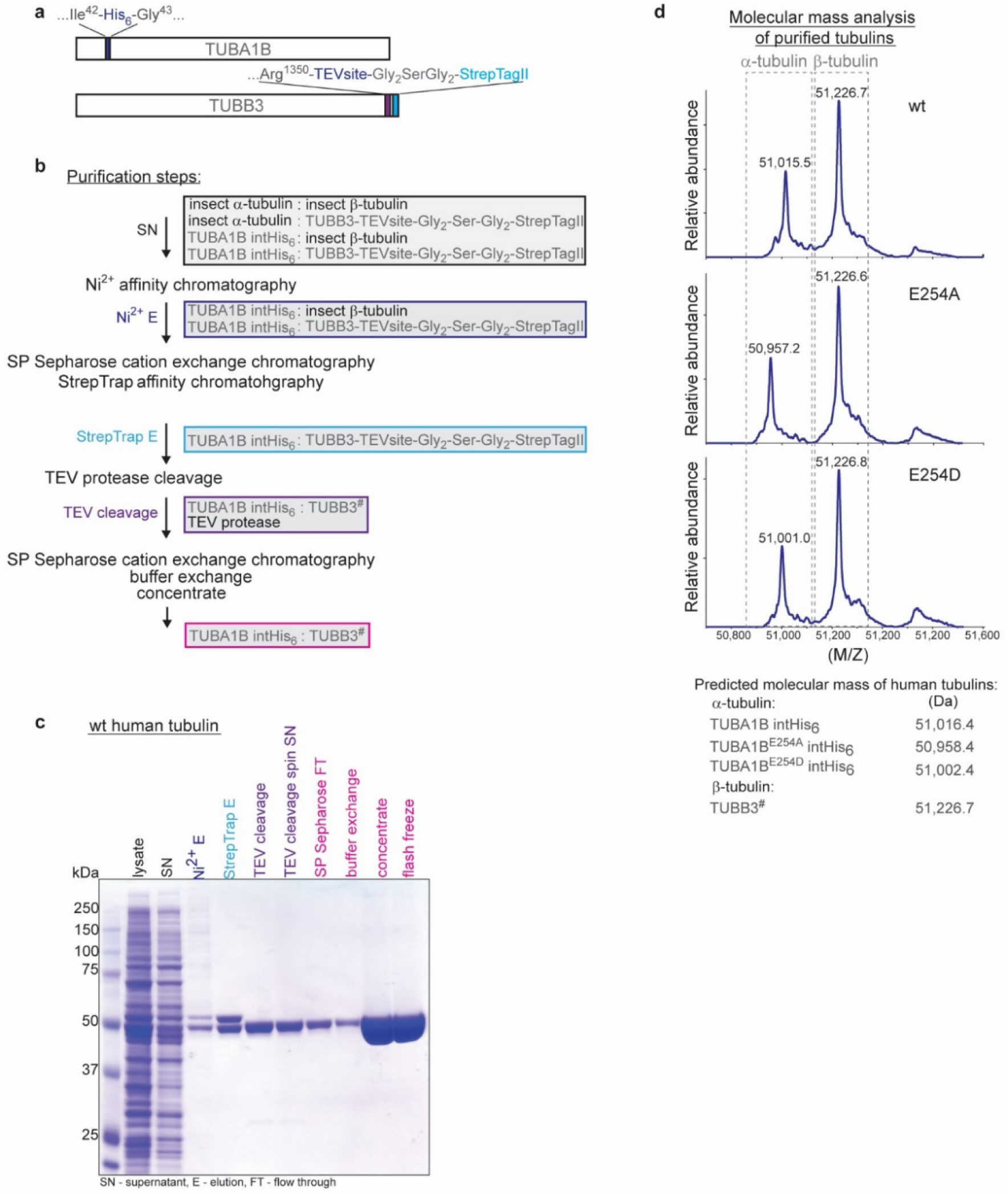
Expression and purification of recombinant human α/β tubulin. **(a)** Schematic of the human TUBA1B α-tubulin and TUBB3 β-tubulin expression constructs. **(b)** Flow chart of the purification steps to obtain recombinant human α/β-tubulin from insect cells. The hash (#) sign on TUBB3 marks the TEV-protease cleavage site. **(c)** Coomassie Blue-stained SDS gel showing the step-wise purification of recombinant wild type (wt) human tubulin. Coloring of the labels as for the corresponding purification steps in the flow chart (b). **(d)** Molecular mass determination of purified recombinant tubulin subunits confirms the lack of insect cell tubulin contamination and uncleaved TUBB3. Note that the predicted masses for TUBA1B and its mutants correspond to N-terminally acetylated versions of the protein. This is a common modification of proteins expressed in insect cells.

**Supplementary Fig. 2:**
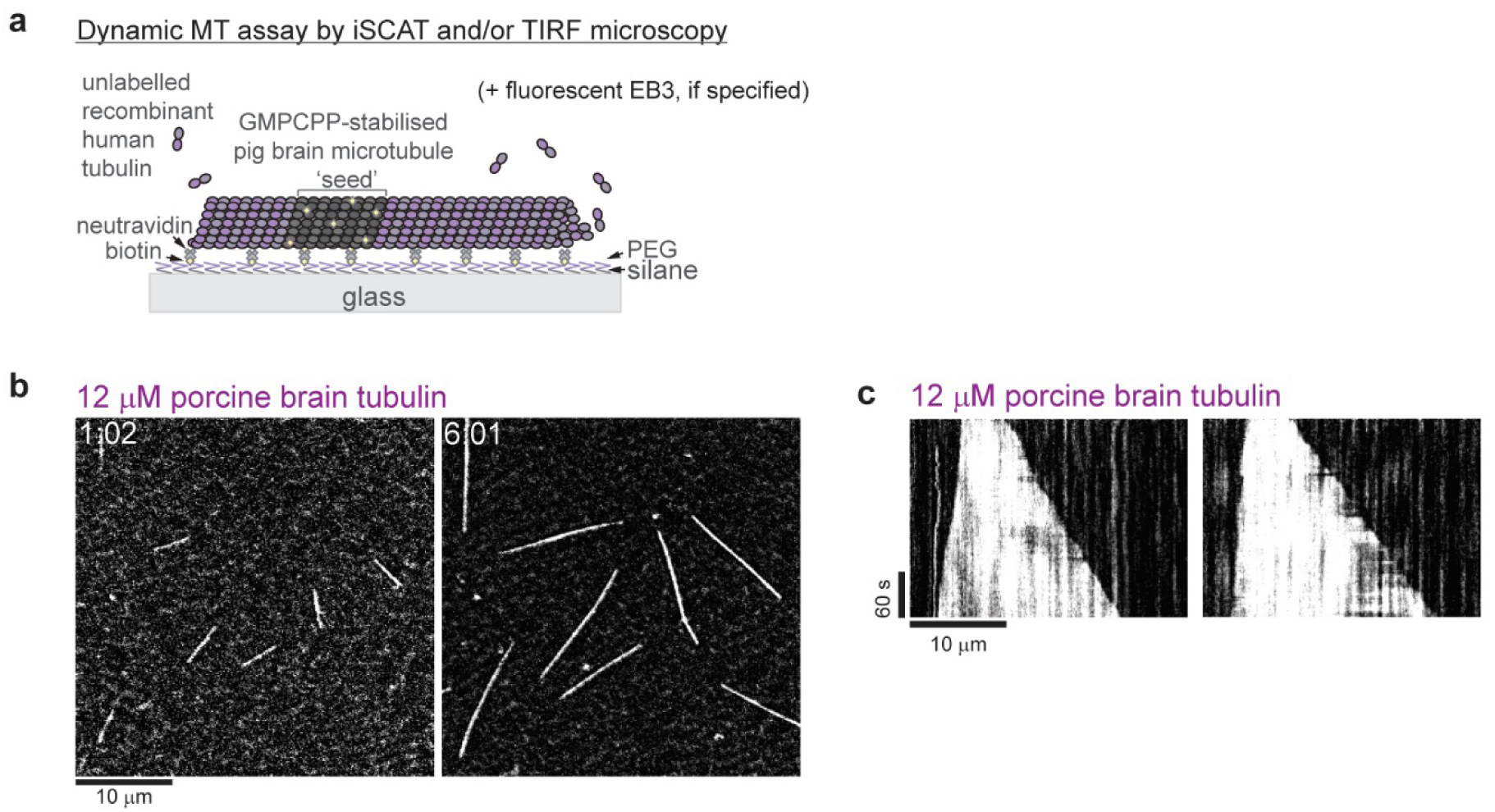
iSCAT microscopy of unlabeled porcine brain microtubules. **(a)** Schematic of a dynamic microtubule assay. **(b)** iSCAT microscopy images of unlabeled porcine brain microtubules growing from GMPCPP-seeds at 12 µM porcine brain tubulin. **(c)** Kymographs showing the characteristic fast plus and slow minus end growth of dynamic porcine brain microtubules. Conditions as in (b). Scale bars as indicated, time is in min:sec.

**Supplementary Fig. 3:**
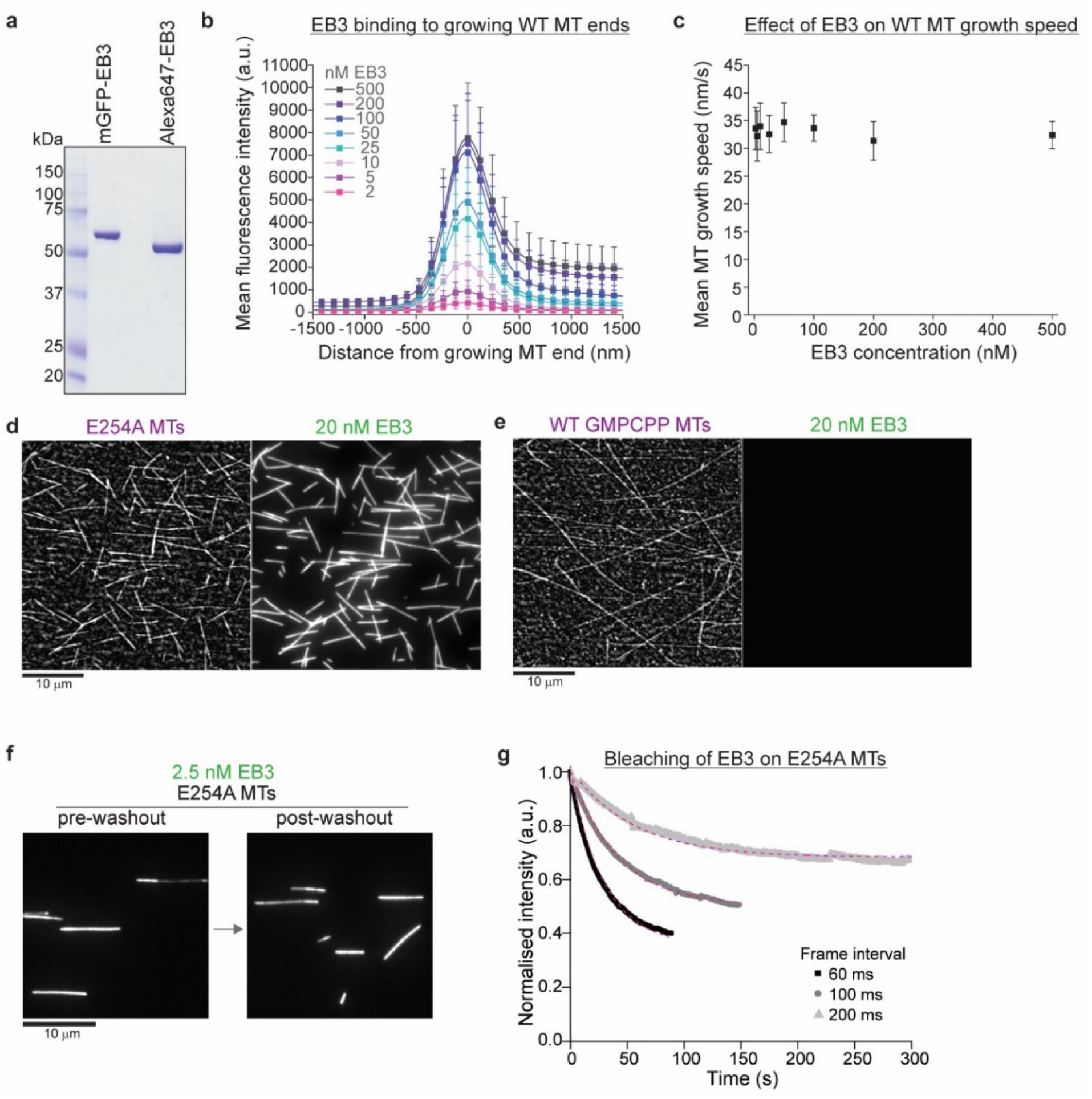
Characterization of EB3 binding to wt and GTPase deficient human microtubules. **(a)** Coomassie Blue-stained SDS gel of purified recombinant mGFP-EB3 and Alexa647-labeled SNAP-EB3 (Alexa647-EB3). **(b)** Mean mGFP-EB3 intensity profiles at the ends of human wt microtubules growing at 12 µM wt tubulin in the presence of varying mGFP-EB3 concentrations (as indicated). Source data as in Fig. 2c. Errors are SD. **(c)** mGFP-EB3 does not affect significantly the growth speed of human wt microtubules. Source data as in (B) and Fig. 2c. **(d, e)** Comparative iSCAT/TIRF microscopy images of 20 nM mGFP-EB3 (green) binding to **(d)** unlabeled E254A microtubules (magenta) and **(e)** wt microtubules polymerized in the presence of GMPCPP, microtubules attached to a Kin1^rigor^ surface. EB3 binds strongly to E254A microtubules, but not to GMPCPP microtubules. **(f)** EB3 washout experiment: TIRF microscopy images of mGFP-EB3 at 2.5 nM bound to unlabeled E254A microtubules immobilized on a Kin1^rigor^ surface before (left) and after (right) washing out mGFP-EB3 from the flow chamber. (**g)** Steady-state bleaching curves for 2.5 nM mGFP-EB3 bound to Kin1^rigor^ surface-immobilized E254A microtubules, imaged with a 60 ms exposure time and different frame intervals, as indicated. Dashed magenta lines are a fit to the data (Methods) revealing a dwell time of mGFP-EB3 molecules of 83 s (and bleaching times of 45 s, 83 s, and 171 s for imaging with a time lapse of 60 ms (single molecule imaging condition), 100 ms, and 200 ms, respectively). Scale bars as indicated, time is in min:sec.

**Supplementary Fig. 4:**
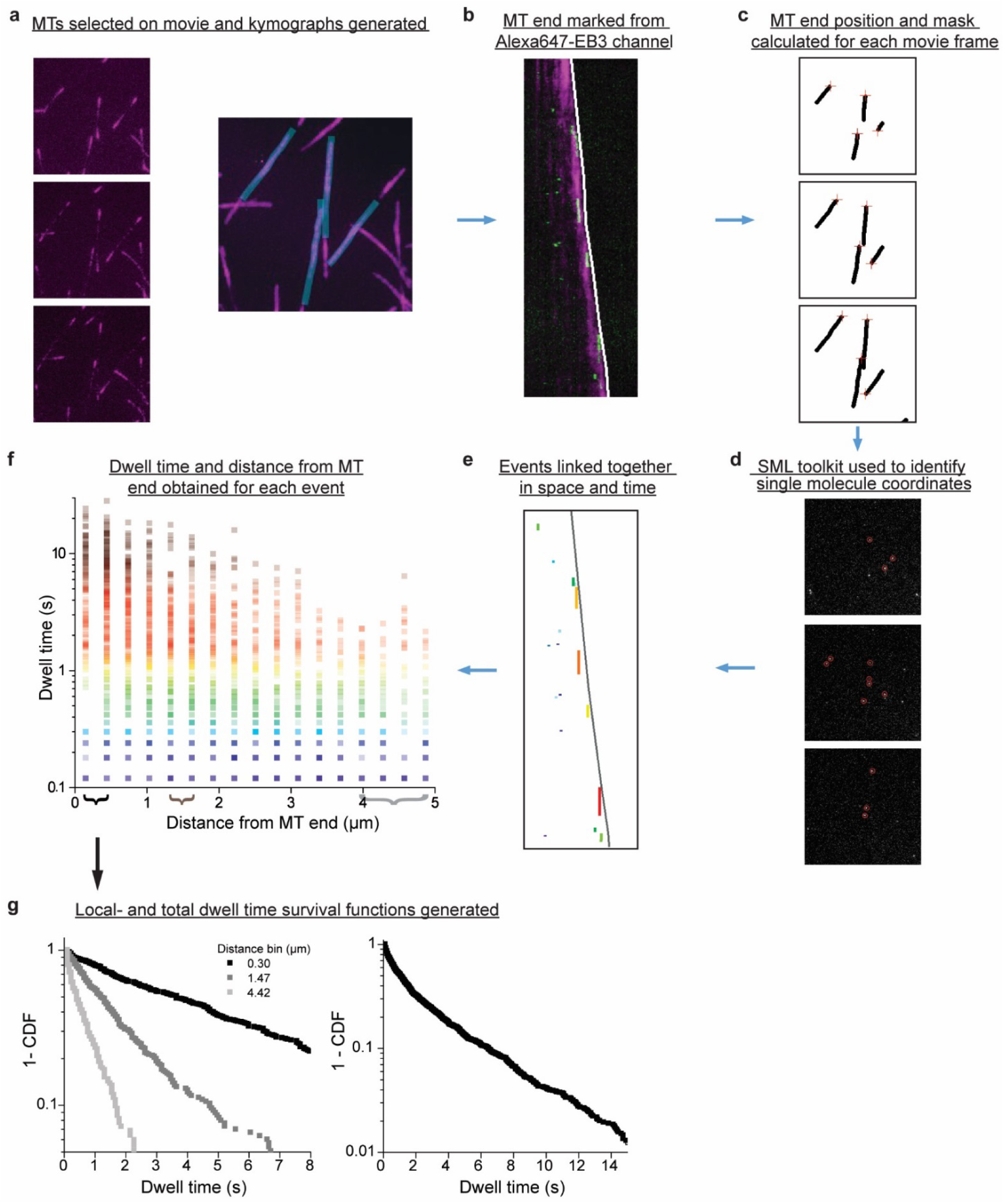
Flow chart of single molecule localization and spatially-resolved dwell time analysis. For single molecule analysis of mGFP-EB3 binding to microtubules, TIRF microscopy movies were recorded in the presence of 25 pM mGFP-EB3 and additional 1 nM Alexa647-EB3 to visualize the growing microtubule end region. **(a**) Left: Single TIRF microscopy frames of a movie showing the Alexa647-EB3 channel. Right: Microtubule positions were marked by hand on a maximum-intensity projection of all frames. (**b)** Kymographs were generated for each marked microtubule in (a), and the growing plus-end position traced by hand in the kymographs (white line). (**c)** The x-y position of each microtubule end was calculated for each frame in the original movie (red crosses). The moving end position and the positions of the static microtubule projection from (a) were used to create a dynamic binary mask movie. (**d)** For each frame of the original movie, the mGFP-EB3 channel was analyzed using a Single Molecule Localization plugin, to determine the coordinates of each potential single molecule event (red circles). The mask from (c) was then used to exclude all events from outside the microtubules of interest. **(e)** Events were linked together in time and space using specific linking parameters. The resulting kymograph corresponding to (b) shows automatically detected and linked single molecule events, color-coded by duration. (**f)** For each linked event, the dwell time and initial distance from the nearest microtubule end were calculated. (**g)** Data from all movies were binned at specific distances from the growing microtubule end (braces in (f)), to give spatially-resolved dwell time survival functions (1-CDF, cumulative density function), or binned over all distances to give a total survival function.

**Supplementary Fig. 5:**
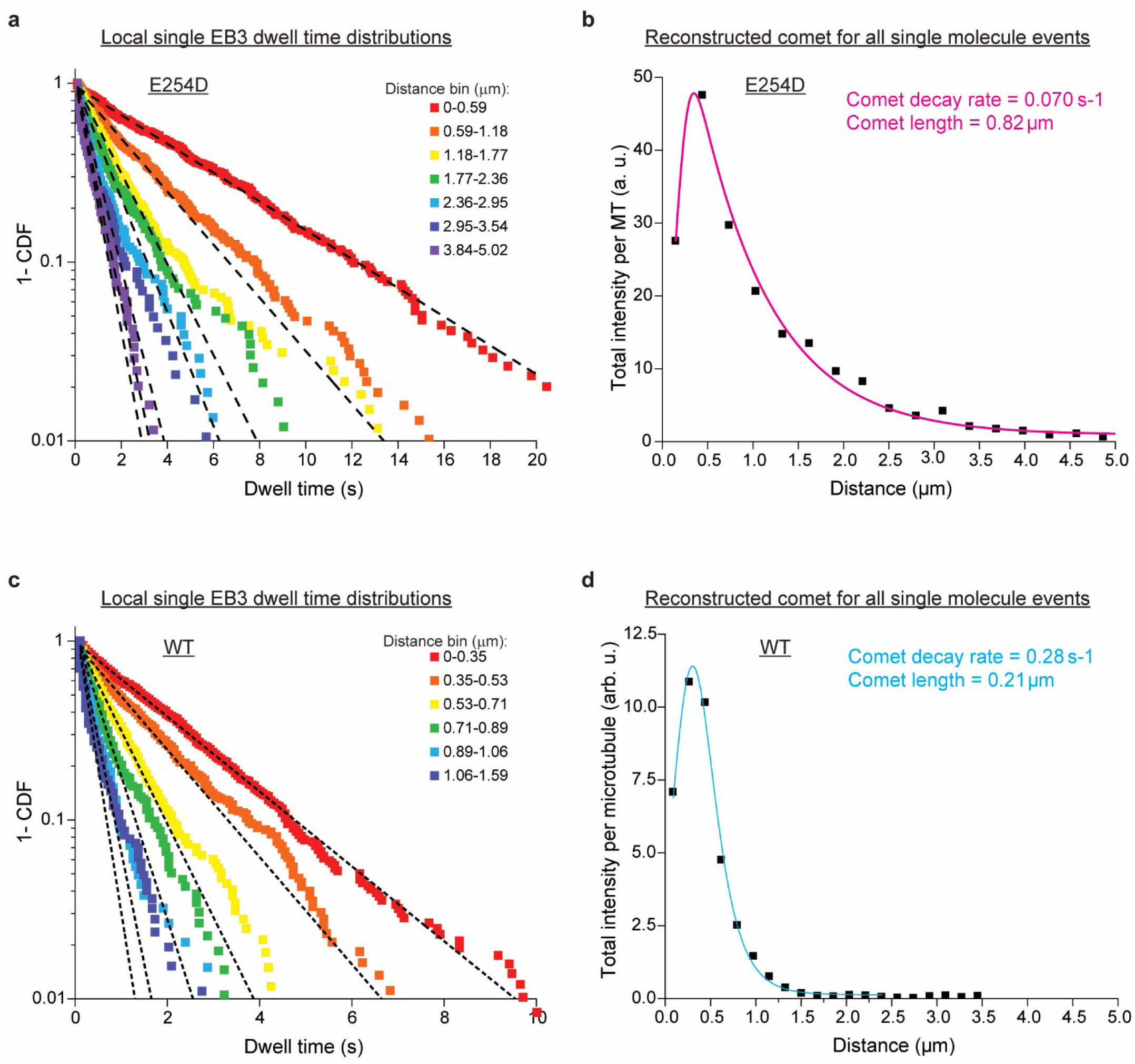
Spatially-resolved EB3 dwell time distributions at the ends of E243D and wt microtubules. **(a)** Local dwell time survival functions at specific distance bins from the microtubule end for E254D microtubules. Dashed black lines are mono-exponential fits. (**b)** Reconstructed comet for all single molecule events, from summing the complete emission of each binding event in each distance bin from the microtubule end. The total intensity is normalized to the number of microtubules analyzed. **(c)** and **(d)**, as (a) and (b) for wt microtubule ends. Intensities in (b) and (d) can be compared. Good agreement of the reconstructed comets based on single molecule data with the measured comets at higher EB3 concentrations (Fig. 3f, 3g), demonstrate equivalence of single molecule and ensemble analysis.

**Supplementary Fig. 6:**
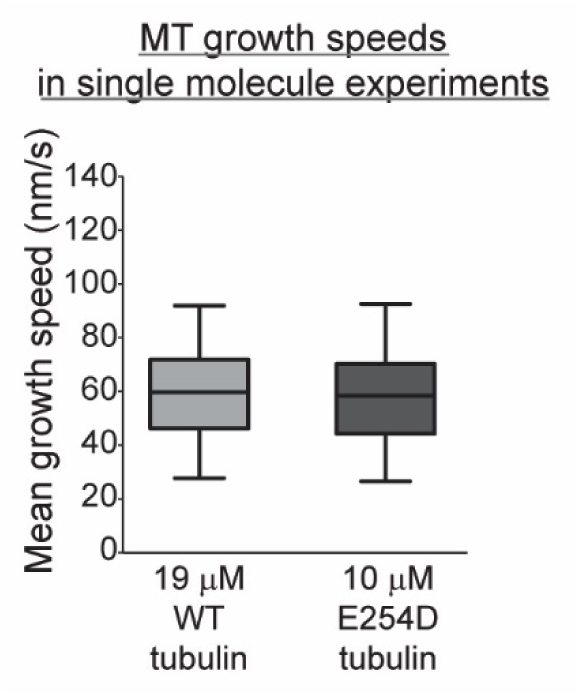
wt and E254D microtubules grow with similar speeds under single molecule experiment conditions. Quantification of microtubule growth speeds at 19 µM wt and 10 µM E254D tubulin concentrations in the presence of 1 nM Alexa647-EB3 and 25 pM mGFP-EB3. The boxes extend from 25^th^ to 75^th^ percentiles, the whiskers extend from 5^th^ to 95^th^ percentiles, and the mean value is plotted as a line in the middle of the box. The source data is the same as presented in Fig. 4.

## REFERENCES

1. Mitchison, T. & Kirschner, M. Dynamic instability of microtubule growth. Nature 312, 237–242 (1984).

2. Brouhard, G.J. & Rice, L.M. Microtubule dynamics: an interplay of biochemistry and mechanics. Nat Rev Mol Cell Biol 19, 451–463 (2018).

3. Carlier, M.F. Guanosine-5’-triphosphate hydrolysis and tubulin polymerization. Review article. Mol Cell Biochem 47, 97–113 (1982).

4. Erickson, H.P. & O’Brien, E.T. Microtubule dynamic instability and GTP hydrolysis. Annu Rev Biophys Biomol Struct 21, 145–166 (1992).

5. Howard, J. & Hyman, A.A. Growth, fluctuation and switching at microtubule plus ends. Nat Rev Mol Cell Biol 10, 569–574 (2009).

6. Akhmanova, A. & Steinmetz, M.O. Control of microtubule organization and dynamics: two ends in the limelight. Nat Rev Mol Cell Biol (2015).

7. Kueh, H.Y. & Mitchison, T.J. Structural plasticity in actin and tubulin polymer dynamics. Science 325, 960–963 (2009).

8. Alushin, G.M. et al. High-resolution microtubule structures reveal the structural transitions in alphabeta-tubulin upon GTP hydrolysis. Cell 157, 1117–1129 (2014).

9. Zhang, R., Alushin, G.M., Brown, A. & Nogales, E. Mechanistic Origin of Microtubule Dynamic Instability and Its Modulation by EB Proteins. Cell 162, 849–859 (2015).

10. Zhang, R., LaFrance, B. & Nogales, E. Separating the effects of nucleotide and EB binding on microtubule structure. Proc Natl Acad Sci U S A 115, E6191–E6200 (2018).

11. Manka, S.W. & Moores, C.A. The role of tubulin-tubulin lattice contacts in the mechanism of microtubule dynamic instability. Nat Struct Mol Biol 25, 607–615 (2018).

12. Bieling, P. et al. CLIP-170 tracks growing microtubule ends by dynamically recognizing composite EB1/tubulin-binding sites. The Journal of Cell Biology 183, 1223–1233 (2008).

13. Bieling, P. et al. Reconstitution of a microtubule plus-end tracking system in vitro. Nature 450, 1100–1105 (2007).

14. Dixit, R. et al. Microtubule plus-end tracking by CLIP-170 requires EB1. Proc Natl Acad Sci U S A 106, 492–497 (2009).

15. Duellberg, C., Cade, N.I., Holmes, D. & Surrey, T. The size of the EB cap determines instantaneous microtubule stability. Elife 5 (2016).

16. Komarova, Y. et al. Mammalian end binding proteins control persistent microtubule growth. The Journal of Cell Biology 184, 691–706 (2009).

17. Maurer, S.P., Fourniol, F.J., Bohner, G., Moores, C.A. & Surrey, T. EBs recognize a nucleotide-dependent structural cap at growing microtubule ends. Cell 149, 371–382 (2012).

18. Roth, D., Fitton, B.P., Chmel, N.P., Wasiluk, N. & Straube, A. Spatial positioning of EB family proteins at microtubule tips involves distinct nucleotide-dependent binding properties. J Cell Sci 132 (2018).

19. Maurer, S.P., Bieling, P., Cope, J., Hoenger, A. & Surrey, T. GTPgammaS microtubules mimic the growing microtubule end structure recognized by end-binding proteins (EBs). Proc Natl Acad Sci U S A 108, 3988–3993 (2011).

20. Nogales, E., Downing, K.H., Amos, L.A. & Lowe, J. Tubulin and FtsZ form a distinct family of GTPases. Nat Struct Biol 5, 451–458 (1998).

21. Anders, K.R. & Botstein, D. Dominant-lethal alpha-tubulin mutants defective in microtubule depolymerization in yeast. Mol Biol Cell 12, 3973–3986 (2001).

22. Kobayashi, T. Dephosphorylation of tubulin-bound guanosine triphosphate during microtubule assembly. J Biochem 77, 1193–1197 (1975).

23. Vemu, A., Atherton, J., Spector, J.O., Moores, C.A. & Roll-Mecak, A. Tubulin isoform composition tunes microtubule dynamics. Mol Biol Cell 28, 3564–3572 (2017).

24. Vemu, A. et al. Structure and Dynamics of Single-isoform Recombinant Neuronal Human Tubulin. J Biol Chem 291, 12907–12915 (2016).

25. Hyman, A.A., Salser, S., Drechsel, D.N., Unwin, N. & Mitchison, T.J. Role of GTP hydrolysis in microtubule dynamics: information from a slowly hydrolyzable analogue, GMPCPP. Mol Biol Cell 3, 1155–1167 (1992).

26. Maurer, S.P. et al. EB1 accelerates two conformational transitions important for microtubule maturation and dynamics. Curr Biol 24, 372–384 (2014).

27. Aher, A. et al. CLASP Suppresses Microtubule Catastrophes through a Single TOG Domain. Dev Cell 46, 40–58 e48 (2018).

28. Doodhi, H. et al. Termination of Protofilament Elongation by Eribulin Induces Lattice Defects that Promote Microtubule Catastrophes. Curr Biol 26, 1713–1721 (2016).

29. Igaev, M. & Grubmuller, H. Microtubule assembly governed by tubulin allosteric gain in flexibility and lattice induced fit. Elife 7 (2018).

30. von Loeffelholz, O. et al. Nucleotide- and Mal3-dependent changes in fission yeast microtubules suggest a structural plasticity view of dynamics. Nat Commun 8, 2110 (2017).

31. Cross, R.A. Microtubule lattice plasticity. Curr Opin Cell Biol 56, 88–93 (2019).

32. Eshun-Wilson, L. et al. Effects of alpha-tubulin acetylation on microtubule structure and stability. Proc Natl Acad Sci U S A 116, 10366–10371 (2019).

33. Ti, S.C., Alushin, G.M. & Kapoor, T.M. Human beta-Tubulin Isotypes Can Regulate Microtubule Protofilament Number and Stability. Dev Cell 47, 175–190 e175 (2018).

34. Geyer, E.A. et al. A mutation uncouples the tubulin conformational and GTPase cycles, revealing allosteric control of microtubule dynamics. Elife 4, e10113 (2015).

## SUPPLEMENTARY REFERENCES

1. Ti, S.C., Alushin, G.M. & Kapoor, T.M. Human beta-Tubulin Isotypes Can Regulate Microtubule Protofilament Number and Stability. Dev Cell 47, 175–190 e175 (2018).

2. Sirajuddin, M., Rice, L.M. & Vale, R.D. Regulation of microtubule motors by tubulin isotypes and post-translational modifications. Nat Cell Biol 16, 335–344 (2014).

3. Vemu, A. et al. Structure and Dynamics of Single-isoform Recombinant Neuronal Human Tubulin. J Biol Chem 291, 12907–12915 (2016).

4. Montenegro Gouveia, S. et al. In vitro reconstitution of the functional interplay between MCAK and EB3 at microtubule plus ends. Curr Biol 20, 1717–1722 (2010).

5. Snapp, E.L. et al. Formation of stacked ER cisternae by low affinity protein interactions. J Cell Biol 163, 257–269 (2003).

6. Zacharias, D.A., Violin, J.D., Newton, A.C. & Tsien, R.Y. Partitioning of lipid-modified monomeric GFPs into membrane microdomains of live cells. Science 296, 913–916 (2002).

7. Jha, R., Roostalu, J., Cade, N.I., Trokter, M. & Surrey, T. Combinatorial regulation of the balance between dynein microtubule end accumulation and initiation of directed motility. EMBO J 36, 3387–3404 (2017).

8. Wasilko, D.J. et al. The titerless infected-cells preservation and scale-up (TIPS) method for large-scale production of NO-sensitive human soluble guanylate cyclase (sGC) from insect cells infected with recombinant baculovirus. Protein Expr Purif 65, 122–132 (2009).

9. Roostalu, J., Cade, N.I. & Surrey, T. Complementary activities of TPX2 and chTOG constitute an efficient importin-regulated microtubule nucleation module. Nat Cell Biol 17, 1422–1434 (2015).

10. Castoldi, M. & Popov, A.V. Purification of brain tubulin through two cycles of polymerization-depolymerization in a high-molarity buffer. Protein Expr Purif 32, 83–88 (2003).

11. Hyman, A. et al. Preparation of modified tubulins. Methods Enzymol 196, 478–485 (1991).

12. Bieling, P., Telley, I.A., Hentrich, C., Piehler, J. & Surrey, T. Fluorescence Microscopy Assays on Chemically Functionalized Surfaces for Quantitative Imaging of Microtubule, Motor, and +TIP Dynamics. Methods Cell Biology 95, 555–580 (2010).

13. Ortega Arroyo, J., Cole, D. & Kukura, P. Interferometric scattering microscopy and its combination with single-molecule fluorescence imaging. Nat Protoc 11, 617–633 (2016).

14. Maurer, S.P. et al. EB1 accelerates two conformational transitions important for microtubule maturation and dynamics. Curr Biol 24, 372–384 (2014).

15. Zhang, R., Roostalu, J., Surrey, T. & Nogales, E. Structural insight into TPX2-stimulated microtubule assembly. Elife 6 (2017).

16. Gebhardt, J.C. et al. Single-molecule imaging of transcription factor binding to DNA in live mammalian cells. Nat Methods 10, 421–426 (2013).

17. Bieling, P. et al. CLIP-170 tracks growing microtubule ends by dynamically recognizing composite EB1/tubulin-binding sites. The Journal of Cell Biology 183, 1223–1233 (2008).

18. Bieling, P. et al. Reconstitution of a microtubule plus-end tracking system in vitro. Nature 450, 1100–1105 (2007).

19. Dixit, R. et al. Microtubule plus-end tracking by CLIP-170 requires EB1. Proc Natl Acad Sci U S A 106, 492–497 (2009).

20. Maurer, S.P., Bieling, P., Cope, J., Hoenger, A. & Surrey, T. GTPgammaS microtubules mimic the growing microtubule end structure recognized by end-binding proteins (EBs). Proc Natl Acad Sci U S A 108, 3988–3993 (2011).

21. Roth, D., Fitton, B.P., Chmel, N.P., Wasiluk, N. & Straube, A. Spatial positioning of EB family proteins at microtubule tips involves distinct nucleotide-dependent binding properties. J Cell Sci 132 (2018).

22. Rickman, J., Duellberg, C., Cade, N.I., Griffin, L.D. & Surrey, T. Steady-state EB cap size fluctuations are determined by stochastic microtubule growth and maturation. Proc Natl Acad Sci U S A 114, 3427–3432 (2017).

24. Bohner, G. et al. Important factors determining the nanoscale tracking precision of dynamic microtubule ends. J Microsc (2015).

25. Ruhnow, F., Zwicker, D. & Diez, S. Tracking single particles and elongated filaments with nanometer precision. Biophys J 100, 2820–2828 (2011).

26. McIntosh, J.R. et al. Microtubules grow by the addition of bent guanosine triphosphate tubulin to the tips of curved protofilaments. J Cell Biol 217, 2691–2708 (2018).

27. Zhang, R., Alushin, G.M., Brown, A. & Nogales, E. Mechanistic Origin of Microtubule Dynamic Instability and Its Modulation by EB Proteins. Cell 162, 849–859 (2015).

28. Hyman, A.A., Salser, S., Drechsel, D.N., Unwin, N. & Mitchison, T.J. Role of GTP hydrolysis in microtubule dynamics: information from a slowly hydrolyzable analogue, GMPCPP. Mol Biol Cell 3, 1155–1167 (1992).

29. Alushin, G.M. et al. High-resolution microtubule structures reveal the structural transitions in alphabeta-tubulin upon GTP hydrolysis. Cell 157, 1117–1129 (2014).

30. Manka, S.W. & Moores, C.A. The role of tubulin-tubulin lattice contacts in the mechanism of microtubule dynamic instability. Nat Struct Mol Biol 25, 607–615 (2018).

31. Zhang, R., LaFrance, B. & Nogales, E. Separating the effects of nucleotide and EB binding on microtubule structure. Proc Natl Acad Sci U S A 115, E6191–E6200 (2018).

32. von Loeffelholz, O. et al. Nucleotide- and Mal3-dependent changes in fission yeast microtubules suggest a structural plasticity view of dynamics. Nat Commun 8, 2110 (2017).

